# Long-range sequential dependencies precede complex syntactic production in language acquisition

**DOI:** 10.1101/2020.08.19.256792

**Authors:** Tim Sainburg, Anna Mai, Timothy Q Gentner

## Abstract

To convey meaning, human language relies on hierarchically organized, long-range relationships spanning words, phrases, sentences, and discourse. The strength of the relationships between sequentially ordered elements of language (e.g., phonemes, characters, words) decays following a power law as a function of sequential distance. To understand the origins of these relationships, we examined long-range statistical structure in the speech of human children at multiple developmental time points, along with non-linguistic behaviors in humans and phylogenetically distant species. Here we show that adult-like power-law statistical dependencies precede the production of hierarchically-organized linguistic structures, and thus cannot be driven solely by these structures. Moreover, we show that similar long-range relationships occur in diverse non-linguistic behaviors across species. We propose that the hierarchical organization of human language evolved to exploit pre-existing long-range structure present in much larger classes of non-linguistic behavior, and that the cognitive capacity to model long-range hierarchical relationships preceded language evolution. We call this the Statistical Scaffolding Hypothesis for language evolution.

**Significance Statement:** Human language is uniquely characterized by semantically meaningful hierarchical organization, conveying information over long timescales. At the same time, many non-linguistic human and animal behaviors are also often characterized by richly hierarchical organization. Here, we compare the long-timescale statistical dependencies present in language to those present in non-linguistic human and animal behaviors as well as language production throughout childhood. We find adult-like, long-timescale relationships early in language development, before syntax or complex semantics emerge, and we find similar relationships in non-linguistic behaviors like cooking and even housefly movement. These parallels demonstrate that long-range statistical dependencies are not unique to language and suggest a possible evolutionary substrate for the long-range hierarchical structure present in human language.

## 2 Introduction

Since Shannon’s original work characterizing the sequential dependencies present in language, the structure underlying long-range information in language has been the subject of a great deal of interest in linguistics, statistical physics, cognitive science, and psychology [1–20]. Long-range information content refers to the dependencies between discrete elements (e.g., units of spoken or written language) that persist over long sequential distances spanning words, phrases, sentences, and discourse. For example, in Shannon’s original work, participants were given a series of letters from an English text and were asked to predict the letter that would occur next. Using the responses of these participants, Shannon derived an upper bound on the information added by including each preceding letter in the sequence. More recent investigations compute statistical dependencies directly from language corpora using either correlation functions [3, 4, 7, 8, 10, 12, 13] or mutual information (MI) functions [2, 5, 6, 14] between elements in a sequence. In both cases, sequential relationships are calculated as a function of the sequential distance between events. For example, in the sequence *a → b → c → d → e → f*, at a distance of three elements, relationships would be calculated over the pairs *a* and *d, b* and *e*, and *c* and *f*.

On average, as the distance between elements increases, statistical dependencies grow weaker. Across many different sequence types, including phonemes, syllables, and words in both text and speech, the decay of long-range correlations and MI in language follows a power law (Eq. 6) [2–14, 18, 19]. This power-law relationship is thought to derive at least in part from the hierarchical organization of language, and has been variously attributed to human language syntax [5], semantics [3], and discourse structure [4]. To understand the link between hierarchical organization in language and a power-law decay in sequential dependencies, it is helpful to consider both the latent and surface structure of a sequence (Fig. 1). When only the surface structure of a sequence is available, as it is for language corpora, a power-law decay in the MI between sequence elements gives evidence of an underlying hierarchical latent structure. This phenomenon can be demonstrated by comparing the MI between elements in a sequence generated from a hierarchically-structured language model, such as a probabilistic context-free grammar (PCFG), to the MI between elements in a sequence generated by a non-hierarchical model, such as a Markov process (Fig. 1). For sequences generated by a Markov process, the strength of the relationship between elements decays exponentially (Eq. 5) as sequential distance increases [5, 21] (Fig. 1A). In the PCFG model, however, linear distances in the sequence are coupled to logarithmic distances in the latent structure of the hierarchy (Fig. 1B-C). While information continues to decay exponentially as a function of the distance in the latent hierarchy (Fig. 1D), this log-scaling results in a power-law decay when MI is computed over corresponding sequential distances (Fig. 1E).

**Figure 1:**
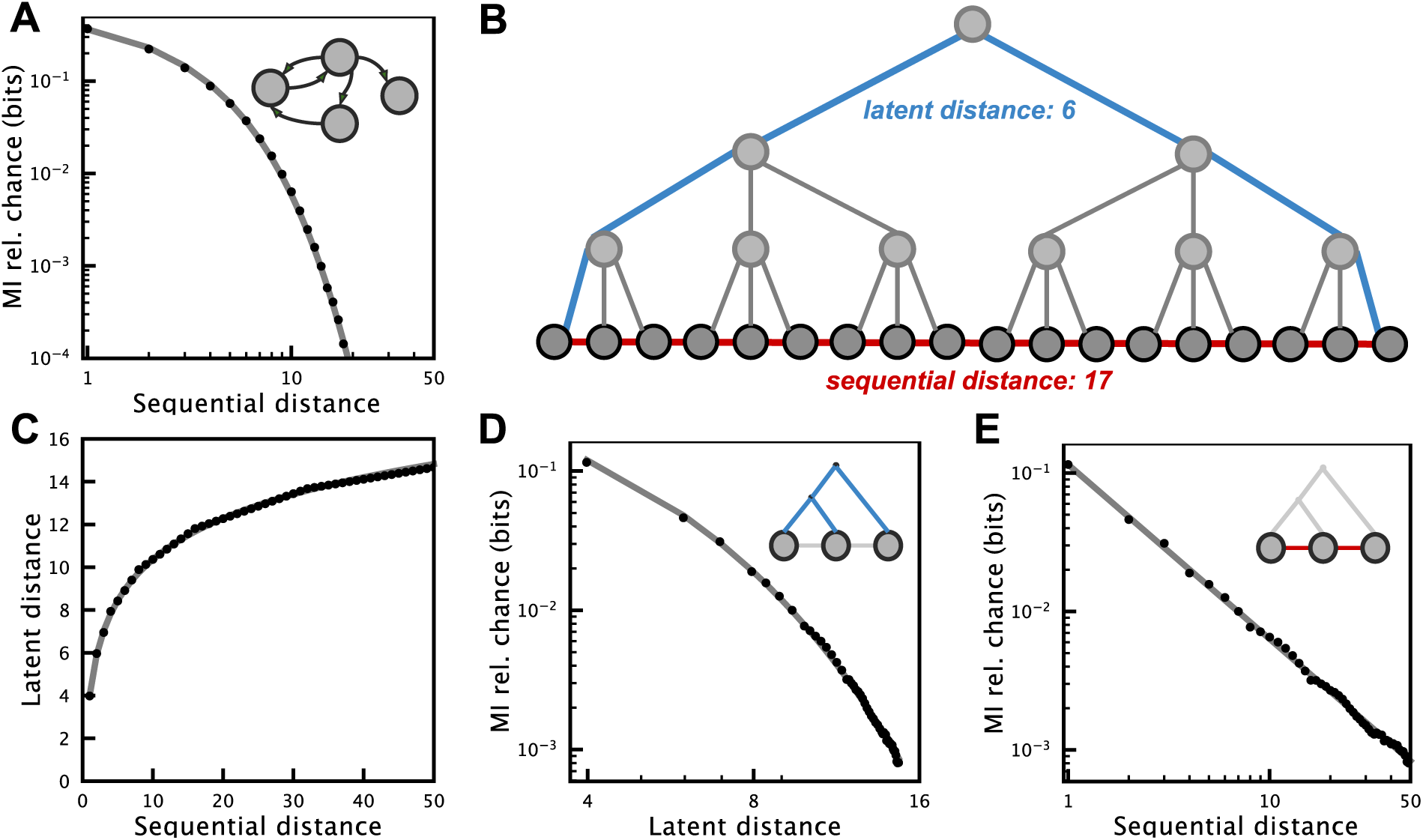
Comparison between sequences with deep latent relationships and iteratively generated sequences. (A) The MI between elements in an iteratively (Markov model) generated sequence decays exponentially as a function of sequential distance. (B) An example sequence with hierarchical latent structure. The latent distance between the two end elements in the sequence is 6 (blue), while the sequential distance is 17 (red). (C) In sequences with hierarchical latent structure, the sequential distance between elements is logarithmically related to the latent distance (fit model: *a ∗ log* _*x∗b*_ + *c* where *x* is sequential distance). (D) Like sequential distance in (A), The MI between elements in a hierarchically generated sequence decays exponentially as a function of latent distance. (E) The MI between elements in a hierarchically generated sequence decays following a power law as a function of sequential distance, which is related to the exponential MI decay seen in (D) and the logarithmic relationship between sequential and latent distance seen in (C). In (A), the probabilistic Markov model used to generate the empirical data has 2 states with a self-transition probability of 0.1. In (C-E) a probabilistic context-free grammar [5] with the same transition probability is used.

In language, long-range relationships convey meaning across hierarchical levels of organization. This latent linguistic structure is thought to underlie the power-law relationships observed across texts and speech [2–5]. The presence of power-law sequential and temporal relationships in natural phenomena is not restricted to human language, however. Here, we demonstrate that the power law underlying long-range statistical relationships in human speech precedes complex morphosyntactic production in language and is part of a larger set of natural behaviors exhibiting similar temporal relationships. The potentially numerous generative mechanisms for these phenomena remain to be established; however their existence evinces a substrate that may have been exploited in the evolution of a cognitive capacity to represent long-range signals prior to the evolution of language.

Beyond language, power-law temporal relationships are observed in both human-unique behaviors like music production [22] and stock market turbulence [23, 24] as well as behaviors that are shared with other animals such as sleep patterns in infants [25] and heart rates in healthy adults [26, 27]. In fact, the ubiquity of power laws in the physical and biological sciences spreads beyond temporal and sequential relationships and is well documented across a variety of phenomena. 1/*f* noise, a power law in the spectral density of a stochastic process, is observed in signals ranging from neural oscillations to flocking patterns in birds [28–31]. The relationship between biological variables often scale following a power law, for example, the allometric scaling laws observed between an organisms size and metabolic rate [32]. A variety of natural distributions such as word frequencies are well described by power-law distributions, a phenomenon termed Zipfs law [33–37]. Power-law distributions are also observed in the connectivity of many biological and social networks, a property called scale-freeness [38–41]. Over much of the past several decades, heated debates have arisen over claims of universal organizing principles of natural phenomena characterized by power laws [28, 31, 34, 41–44].

Across the diverse phenomena described by power-law relationships in the natural sciences, one commonality is that the origins of the observed power law are still not fully understood and mechanistic implications of power laws are often overstated [28, 31, 34, 41, 43, 44]. Although mechanisms have been proposed to account for the various forms of power laws observed in natural phenomena, the presence alone of a power law provides little insight into the underlying generative mechanism [31, 34, 42–44]. This is true of language as well. While the power laws characterized in language are consistent with generative mechanisms posited in syntactic theory [5, 45], they are not confirmatory. The presence of a power law in language does confirm, however, that relationships spanning long distances exist in the signal. Given the presence of power-law sequential relationships in human language, the question remains whether the power law is a product of linguistic structure, or whether these relationships originate in lower-level phenomena that are not unique to human language. If long-range relationships predate the evolution of language, they may have influenced the structure of temporal relationships that evolved with language.

Beyond human language, numerous other human behaviors [46–51], animal behaviors [52–57], animal vocalizations [37, 58–66], and other biologically-generated processes [25–27, 31, 67–70] have been described as being hierarchically organized or display long-timescale organization. Such behaviors range from the seemingly non-complex patterns of behavior exhibited by fruit flies [52, 56] to tool usage in great apes [53, 54]. For this reason, it has been argued that hierarchical organization is an inherent property of biological processes, including human behavior [50, 71, 72] and that the hierarchical structure of behavior is inherited from the lower-level organization of neurophysiological mechanisms that produce it [73–76], which themselves can be characterized by power-law relationships in temporal sequencing [29, 30, 77]. The developmental and/or evolutionary dependence of linguistic structure on underlying, domain-general, cognitive and neural processes has been posited by several researchers [50, 51, 76, 78].

Despite the numerous observations of hierarchical structure and long-range dependencies in non-human animal behaviors, few studies have examined the statistical dynamics of these behaviors quantitatively. Those that do have found power-law dynamics in the communication and behaviors of animals that are phylogenetically distant from humans [2, 79–81]. This, along with the prevalence of long-range power-law relationships in other natural phenomena [28, 31], supports the generality of these organizing principles across all behaviors. On the other hand, sequential organization in the vocal communication signals of non-human primates may extend over only a few elements [82, 83], and descriptions of hierarchical non-vocal behaviors in non-human primates tend to only be a few elements long [53, 54, 84], supporting at most a very shallow hierarchical structure. Thus, the extent to which a power-law decay provides a unified description of long-range statistical dependencies in behavior has yet to be determined. This question has particular relevance to human language, where it is unknown whether power-law relationships in sequential organization are present throughout language development, or emerge as linguistic structure develops. Understanding the ubiquity of power-law relationships across non-linguistic and non-human behavior, as well as across human language acquisition, may help to explain the origins of this organizing principle in language.

### 2.1 Present work

In the present work, we perform three groups of analyses exploring whether non-linguistic and pre-linguistic long-range statistical relationships parallel the long-range statistical relationships present in adult language. First, we analyze a series of language development corpora of children learning English, starting at six months of age [85–98], to determine whether long-range relationships are present in human vocalizations prior to the production of hierarchically-organized linguistic structure. Second, we analyze the long-range statistical dependencies of a human non-linguistic corpus of transcribed actions taken by humans while cooking [99], to determine whether power-law relationships are present in the sequential organization of non-linguistic human behaviors. Finally, we analyze the long-range sequential relationships in datasets of freely moving fruit flies (*Drosophila melanogaster*) [56] and zebrafish (*Danio rerio*) behavior [100], both of which have been previously characterized as being hierarchically organized, to determine whether a power law is present in the sequential organization of non-human non-linguistic behavior.

We show that both human non-linguistic and non-human non-linguistic behavior exhibits long-range power-law statistical dependencies like those observed in mature human language. In child language datasets, we observe a power-law as early as 6 to 12 months of age, while children are still in the “babbling” stage of language development. In the animal behavior datasets, we observe long-range power-law decays spanning many minutes (*>*6 minutes in *Drosophila* and *>*20 minutes in zebrafish).

## 3 Results

### 3.1 Language acquisition

Although much work has explored the information content and long-range sequential organization of human language, relatively few studies have examined these properties in speech [2] or language development directly. Here we investigate the long-range information present in speech during language development using datasets from the TalkBank project [85, 86].

We first examined MI decay in sequences of words over nine datasets of natural speech from English speaking children included in the CHILDES repository [86, 91–98] and three datasets of sequences of phonemes from the PhonBank repository [85, 87–89], both of which are part of the TalkBank repository [86]. Each dataset within CHILDES and PhonBank was collected in a slightly different manner. In our analyses, we included only transcripts of spontaneous speech that were collected from typically-developing children (usually at an in-home setting with family or an experimenter). The subset of CHILDES we used includes word-level transcripts of speech from children aged 12 months to 12 years of age. The subset of PhonBank we used includes phonetic transcriptions of speech given in the International Phonetic Alphabet (IPA) from children aged 6 months to four years of age. Between the phoneme and word-level datasets, a large range of speech and language development is covered.

For the MI analysis on phonemes, we binned transcripts into five 6-month age groups (6-12, 12-18, 18-24, 24-30, 30-36) and one age group from 3 years to 4 years. Each transcript was analyzed as sequences of phonemes, where phoneme distributions for each transcript are treated independently to account for variation in acquired vocabulary across individuals during development. Because transcript lengths varied between age groups (Fig. S1), we analyzed MI at sequential distances up to the median transcript length for each age group. Across all age groups, the decay in MI over sequences of phonemes is best fit by a composite power-law and exponential decay model (Fig. 2A-C; relative probabilities 0.897 to >0.999; Table S2). In each age group, we observe both a clear power law prominent over long distances (Fig. 2B) and a clear exponential decay at short word distances (Fig. 2C), consistent with prior results on adult speech [2].

**Figure 2:**
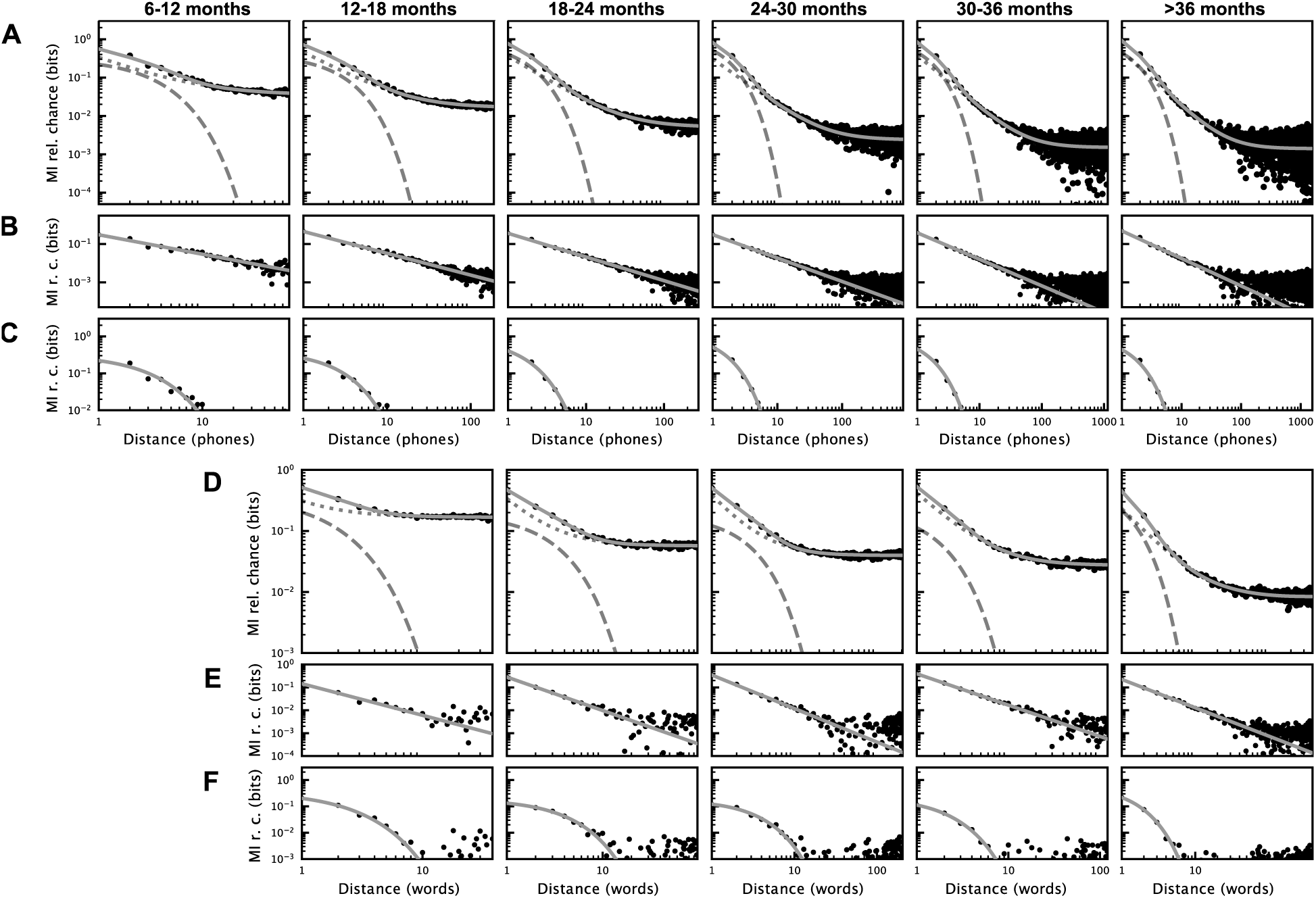
Mutual Information decay over words and phonemes during development. (A) MI decay over phonemes for each age group. MI decay is best fit by a composite model (solid grey line) for all age groups across phonemes and words. Exponential and power-law decays are shown as a dashed and dotted grey lines, respectively. (B) The MI decay (as in (A)) with the exponential component of the fit model subtracted to show the power-law component of the decay. (C) The same as in (B), but with the power-law component subtracted to show exponential component of the decay. (D-F) The same analyses as A-C, but for words.

For the MI analysis on words, we binned transcripts into four 6-month age groups (12-18, 18-24, 24-30, 30-36) and one age group from 3 years to 12 years. The MI decay between words is best fit by a composite model of power-law and exponential decay (Eq. 7; relative probability = 0.989 for 12-18 months and > 0.999 for all other age groups; Fig. 2D-F; Table S1).

We also computed the MI decay over control sequences of words and phonemes that had been shuffled to isolate sequential relationships at different levels of organization (e.g. phoneme, word, utterance, transcript; Figs. S2, S3, S4). Consistent with Sainburg et al., [2], we observe that short-range relationships captured by exponential decay are largely carried within words and utterances, while long-range relationships captured by a power-law decay are carried across longer timescales between words and utterances. In particular, long-range relationships are eliminated when between-utterance structure is removed by randomly shuffling the order of utterances within a transcript (Figs. S2E, S3C) and retained when within-utterance structure is removed by shuffling words or phonemes within utterances (Figs. S2D, S3B) or phonemes within words (Fig. S2C). When MI decay is computed over part-of-speech labels for the words in CHILDES, we find a transition from MI decay that is best fit by a power-law decay alone at 12-24 months of age, to MI decay that is best fit by a composite model of power-law and exponential decay after 24 months (Fig S3D). Shuffling word order eliminates all long-range sequential relationships while preserving short timescale exponential relationships (Figs. S2B, S3E), and shuffling phoneme order within transcripts removes all sequential relationships (Figs. S2F). Across each shuffling analysis, we observe that short-range information content captured by exponential decay is largely captured within words and utterances, while long-range information is carried between utterances, even during early language production.

As an additional control to ensure that the observed MI decay patterns are not the product of mixing datasets from multiple individuals, we also computed the MI decay of the longest individual transcripts comprising each age cohort across both phonemes and words. The decay of the longest individual transcripts parallels the results across transcripts from Fig. 2 (Figs. S5, S6).

### 3.2 Human behavior

To contrast the long-range statistical structure of human language with non-linguistic human behaviors, we require a relatively large dataset of long, discrete, sequences of behavior. We chose the Epic Kitchens dataset [99], as it was the largest available segmented dataset of long sequences of individual actions, and because cooking has previously been described as having complex hierarchical syntactic structure [101].

The Epic Kitchens dataset consists of a series of videos in which each section of the video is labeled with an action and noun, for example *open door → turn-on light → close door → open fridge → …*. We calculate MI only over the sequences of verb classes, of which there are 119 unique classes. We computed the MI up to a distance of the median sequence length of 45 actions.

In contrast with the speech datasets, we found that the Epic Kitchens dataset was best fit by a power-law decay model with no exponential component (Eq. 6; Fig. 3; relative probability = 0.597; Table S3). We additionally looked at the MI decay of the longest cooking transcripts and found the MI decay of individual sequences were similar to MI decay across the entire dataset (Fig S7).

**Figure 3:**
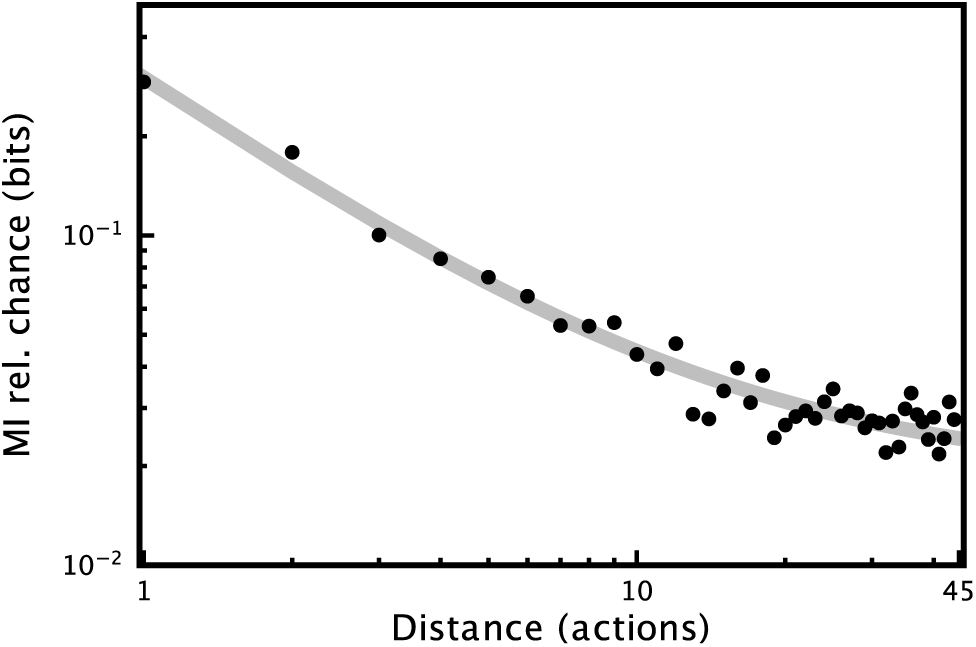
Mutual Information decay over actions in the Epic Kitchens dataset [99]. Data is fit by a power-law decay model (Eq. 6).

### 3.3 Animal behavior

The datasets of animal behavior used in our analyses were videos of zebrafish [100] and *Drosophila* [56] *movements that had been transcribed in an unsupervised manner, i*.*e without external reference to a priori* state labels. In both datasets, raw data recorded from individual animals were projected into a low-dimensional space and were then clustered into discrete states. These states were then labelled *post hoc* with human-interpretable descriptions such as “slow”, “side leg”, or “anterior” for *Drosophila*, and “O-bend” or “J-turn” for zebrafish. *Drosophila* behavior has a long history of being described in hierarchical terms [52, 56, 102], and the dataset used here, in particular, demonstrates long-range relationships extending over hundreds to thousands of states [56]. The zebrafish dataset used here has also previously been shown to contain sequential information that unfolds over multiple timescales [100, 103]. Both datasets were chosen because they contain large sets of discrete behaviors from individuals over long periods of time.

In both the zebrafish and *Drosophila* datasets, we observe an MI decay that is best fit by a composite power-law and exponential decay model (Fig. 4; relative probabilities > 0.999; Table S3). The shape of the MI decay differs somewhat between the two datasets, however. In the case of the zebrafish, the relative contributions of the exponential and power-law components of the decay mirror the results obtained in speech. That is, an exponential component to the decay is observed at short distances under 10 elements, which gives way to a power-law at longer distances. In the case of the *Drosophila*, the power-law component of the decay is dominant throughout the signal, and the exponential component of the decay only captures a small portion of the variance at a distance of around 10-200 elements.

**Figure 4:**
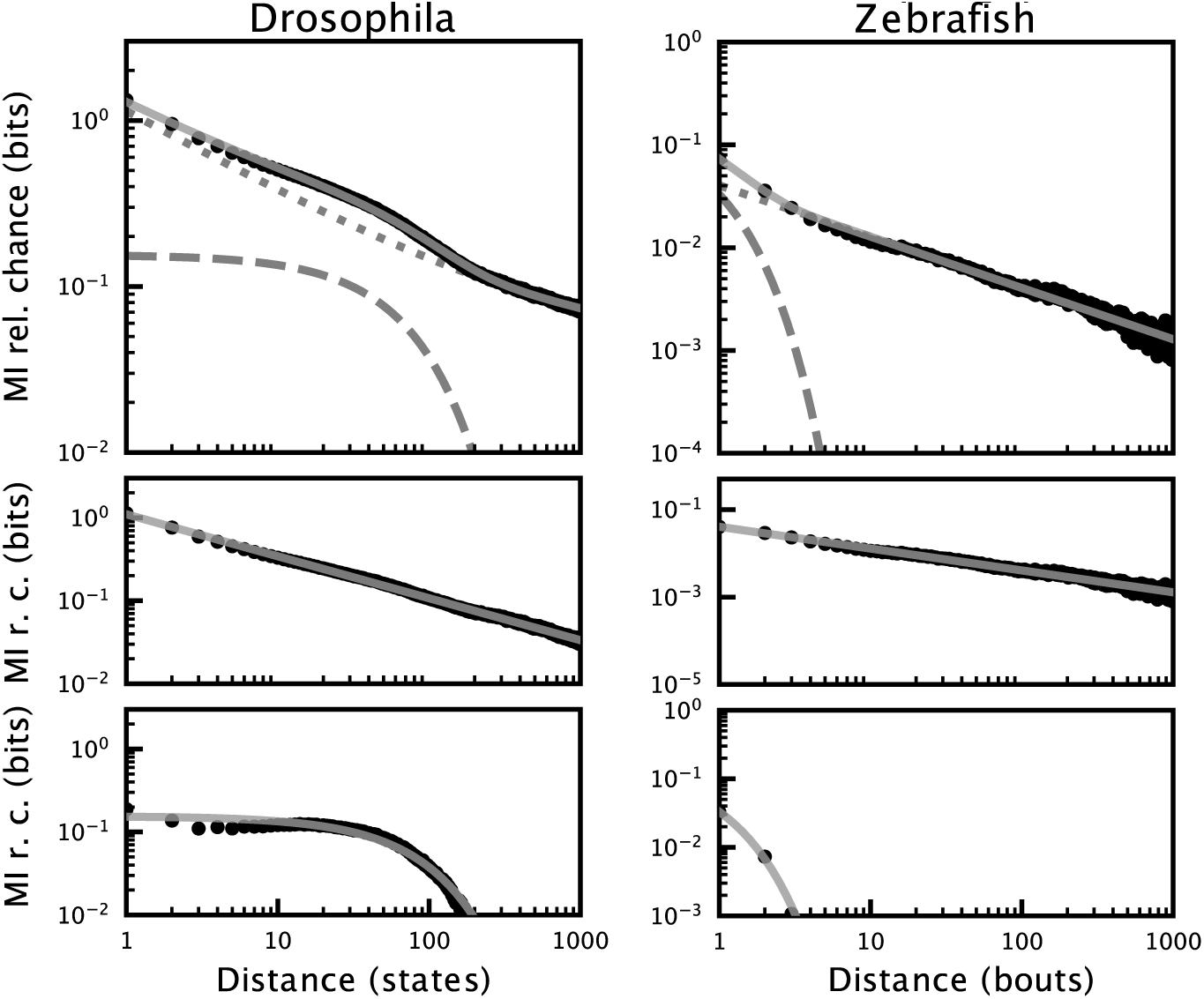
Mutual Information decay over Zebrafish and *Drosophila* behavior. Data is displayed in the same manner as Fig. 2.

We additionally looked at a subset of the longest individual transcripts of *Drosophila* (Fig. S8) and zebrafish (Fig. S9) behavior and found that MI decay at the individual level varies between individual transcripts but matches the long-range decay observed across the datasets.

## 4 Discussion

We analyzed the long-range sequential information present in language production during development, and several sequentially organized and putatively hierarchical non-linguistic behaviors in other species. In all cases, the information between behavioral elements decays following a power law as sequential distance increases. For language, we find that that the long-range statistical relationships characteristic of adult usage [2] are present as early as 6 to 12 months in phonemes and 12-18 months in words, preceding the production of complex linguistic structure [84]. We see similar long-range power-law structure in the sequential organization of human food preparation and cooking. Cooking is a relatively modern and human-unique behavior [104], however, and may have arisen after humans developed more deeply hierarchical and highly planned tool usage behaviors [84, 105]. Yet, we also observe similar long-range organization in the movement patterns of *Drosophila* and zebrafish, consistent with previous reports for birdsong [2]. Long-range statistical relationships are present developmentally in speech before hierarchical linguistic structures are produced, and exist in widely varying animal species. Thus, the long-range statistical relationships present in language are not unique to linguistic behaviors or to humans.

These results compel reconsideration of the mechanisms that shape long-range statistical relationships in human language. Traditionally, the power-law decay in information between the elements of language (phonemes, words, etc.) has been thought to be imposed by the hierarchical linguistic structure of syntax, semantics, and discourse [3–5]. Early development provides a natural experiment in which one can examine human vocal communication absent the production of complex syntactic and semantic structures. Remarkably, even at a very early age, prior to the production of mature syntactic structures, vocal sequences show adult-like long-range dependencies. This does not rule out the possibility that long-range dependencies in adult language are driven in part by linguistic structure, but this hierarchical organization alone cannot explain our observations. What seems most reasonable to us, is that multiple mechanisms impose long-range dependencies on human speech and language, and that these operate on different developmental timescales. We take our observations of similar power laws in diverse non-linguistic behaviors to reinforce the idea that multiple mechanisms impose power-law dynamics on behavioral sequences. Indeed, power-laws are found in natural phenomena as distant from language as the sequential organization of earthquakes [106] and river water levels [107]. It may be that the power-law structure of human language reflects a very deep embedding of multiple, hierarchically structured complex systems, at varying levels of abstraction from linguistic, to motor control, to even more general underlying processes. Understanding the various power-law relationships in natural phenomena, and their origins, remains an area of active research [28, 31, 42].

Regardless of any deeper understanding of underlying mechanisms, our results demonstrate clear patterns in the information conveyed across time in both linguistic and non-linguistic behaviors. These patterns exist. Thus, they are potentially available and useful to any cognitive agent that engages with them. For example, in the movement patterns of a housefly, evolutionary fitness may be conferred to individuals (e.g. predators or mates) that can better anticipate the behavior of others by integrating long-range statistical dependencies. For human language, these selective advantages and abilities seem clear, as sensitivity to long-range organization has obvious benefit for comprehension. Outside of language, evidence for long-range sensitivities is more sparse, but humans do show scale invariance in retrospective memory tasks [108] and attention to power-law timescales in anticipation of future events in cognitive tasks [109]. The extent to which non-human animals are sensitive to the long-range dynamics (power-law or otherwise) of information in the environment is unknown. If non-human animals can model the long-range statistical dependencies present in their environment, this capacity would constitute a component of the broad faculty of language [110], that is, a necessary, but not uniquely-human, component of language. The presence of long-range statistical dependencies in non-linguistic behaviors and a generalized perceptual sensitivity to them would provide a scaffold on which language could evolve, and where hierarchical syntax and semantics can be understood as later additions that exploit existing long-range structures and sensitivities. We refer to this idea as the Statistical Scaffolding Hypothesis.

## 5 Methods

### 5.1 Mutual information

For each dataset, we calculate the sequential MI over the elements of the sequence dataset (e.g. words produced by a child, actions performed by *Drosophila*). Each element in each sequence is treated as unique to that transcript to account for different distributions of behaviors across different transcripts within datasets.

Given a sequence of discrete elements *a → b → c → d → e* We calculate mutual information as:

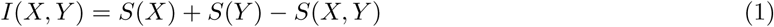

Where *X* and *Y* are the distributions of single elements at a given distance. For example, at a distance of two, *X* is the distribution [*a, b, c*] and *Y* is [*c, d, e*] from the set of element-pairs (*a* − *c, b* − *d*, and *c* − *e*). *Ŝ*(*X*) and *Ŝ*(*Y*) are the marginal entropies of the distributions of *X* and *Y*, respectively, and *Ŝ*(*X, Y*) is the entropy of the joint distribution of *X* and *Y*.

To estimate entropy, we employ the Grassberger [111] method which accounts for under-sampling true entropy from finite samples:

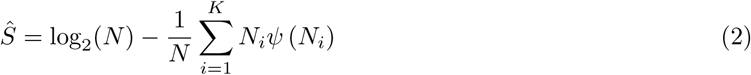

where *ψ* is the digamma function, *K* is the number of categories of elements (e.g. words or phones) and *N* is the total number of elements in each distribution.

We then adjust the estimated MI to account for chance. To do so, we subtract a lower bound estimate of chance MI (*Î*_*sh*_):

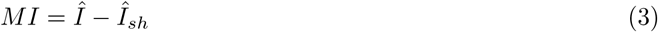

This sets chance MI at zero. We estimate MI at chance (*Î*_*sh*_) by calculating MI on permuted distributions of labels *X* and *Y*:

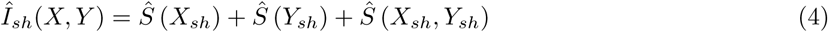

*X*_*sh*_ and *Y*_*sh*_ refer to random permutations of the distributions *X* and *Y* described above. Permuting *X* and *Y* effects the joint entropy *S*(*X*_*sh*_, *Y*_*sh*_) in *I*_*sh*_, but not the marginal entropies *S*(*X*_*sh*_) and *S*(*Y*_*sh*_). *Î*_*sh*_ is related to the Expected Mutual Information [112–114] which accounts for chance using a hypergeometric model of randomness.

Importantly, MI calculated over a sequence as a function of distance is referred to as a “mutual information function”, to distinguish it as the functional form of mutual information, which measures the dependency between two random variables [14]. In the mutual information function, samples from the distributions *X* and *Y* are taken from the same sequence, thus they are not independent. MI as a function of distance acts as a generalized form of the correlation function that can be computed over symbolic sequences and captures non-linear relationships [14].

### 5.2 Fitting mutual information decay

We fit the three following models:

An exponential decay model:

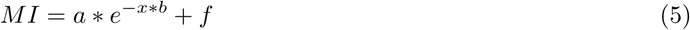

A power-law model:

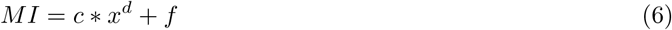

A composite model of the power-law and exponential models:

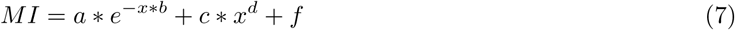

where *x* represents the inter-element distance between units (e.g. phones or syllables).

To fit the model on a logarithmic scale, we computed the residuals between the log of the MI and the log of the models estimation of the MI. We scaled the residuals during fitting by the log of the distance between elements to emphasize fitting the decay in log-scale because distance was necessarily sampled linearly as integers. Models were fit using the lmfit Python package [115] using Nelder-Mead minimization. We compared model fits on the basis of AICc and report the relative probability of each model fit to the MI decay [2, 116]. The parameters for each best-fit model for Figs 2, 3, and 4 can be found in Table 4.

**Table 1:** CHILDES dataset model fit results for each decay model as shown in Fig. 2.

|  | 12-18 months | 18-24 months | 24-30 months | 30-36 months | 3+ years |
| --- | --- | --- | --- | --- | --- |
| **AICc** |  |  |  |  |  |
| exp. | -313.876 | -696.201 | -1464.82 | -735.697 | -2314.6 |
| combined | -322.789 | -819.742 | -1737.37 | -951.049 | -2989.21 |
| power-law | -296.67 | -746.061 | -1623 | -933.579 | -2939.72 |
| $r^2$ | | | | | |
| exp. | 0.997 | 0.992 | 0.991 | 0.986 | 0.973 |
| combined | 0.998 | 0.998 | 0.998 | 0.998 | 0.995 |
| power-law | 0.995 | 0.995 | 0.996 | 0.997 | 0.994 |
| **Relative likelihood** |  |  |  |  |  |
| exp. | 0.012 | <0.001 | <0.001 | <0.001 | <0.001 |
| combined | >0.999 | >0.999 | >0.999 | >0.999 | >0.999 |
| power-law | <0.001 | <0.001 | <0.001 | <0.001 | <0.001 |
| **Relative probability** |  |  |  |  |  |
| exp. | 0.011 | <0.001 | <0.001 | <0.001 | <0.001 |
| combined | 0.989 | >0.999 | >0.999 | >0.999 | >0.999 |
| power-law | <0.001 | <0.001 | <0.001 | <0.001 | <0.001 |

**Table 2:** PhonBank dataset model fit results for each decay model as shown in Fig. 2.

|  |  | 6-12 months | 12-18 months | 18-24 months | 24-30 months | 30-36 months | 3+ years |
| --- | --- | --- | --- | --- | --- | --- | --- |
| **AICc** | **exp.** | -5687.13 | -4842.29 | -4240.44 | -1371.61 | -1091.13 | -417.621 |
|  | **combined** | -5998.03 | -5302.25 | -5025.96 | -1903.5 | -1522.77 | -484.369 |
|  | **power-law** | -5993.7 | -5288.72 | -4971.83 | -1836.16 | -1369.5 | -437.315 |
| $r^2$ | **exp.** | 0.803 | 0.878 | 0.928 | 0.967 | 0.983 | 0.989 |
|  | **combined** | 0.841 | 0.919 | 0.971 | 0.995 | 0.998 | 0.996 |
|  | **power-law** | 0.841 | 0.918 | 0.969 | 0.994 | 0.996 | 0.992 |
| **Relative likelihood** | **exp.** | <0.001 | <0.001 | <0.001 | <0.001 | <0.001 | <0.001 |
|  | **combined** | >0.999 | >0.999 | >0.999 | >0.999 | >0.999 | >0.999 |
|  | **power-law** | 0.115 | 0.001 | <0.001 | <0.001 | <0.001 | <0.001 |
| **Relative probability** | **exp.** | <0.001 | <0.001 | <0.001 | <0.001 | <0.001 | <0.001 |
|  | **combined** | 0.897 | 0.999 | >0.999 | >0.999 | >0.999 | >0.999 |
|  | **power-law** | 0.103 | 0.001 | <0.001 | <0.001 | <0.001 | <0.001 |

**Table 3:**
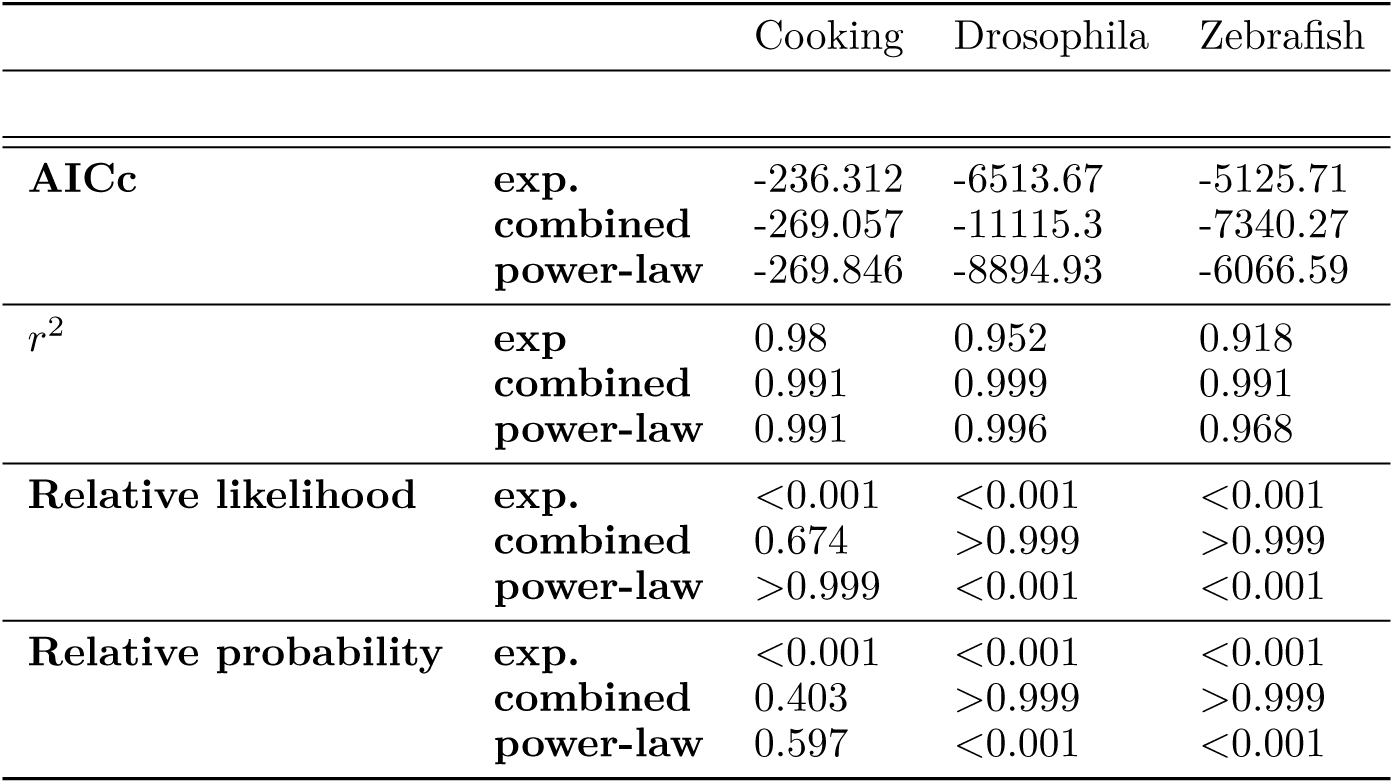
Epic Kitchens, Drosophila, and Zebrafish model fit results at 45, 1000, and 1000 elements of distance respectively.

**Table 4:** MI decay parameters for Figs 2, 3, and 4. The parameters correspond to Equation 7 (*a e*^−*x∗b*^ + *c * x*^*d*^ + *f*). *a* and *b* for the Cooking dataset are not shown because the best-fit model is the power-law model.

| Dataset | Age (yrs) | $a$ | $b$ | $c$ | $d$ | $f$ |
| --- | --- | --- | --- | --- | --- | --- |
| CHILDES | 1-1.5 | 0.387±0.101 | 0.645±0.113 | 0.145±0.038 | -1.382±0.345 | 0.168±0.003 |
|  | 1.5-2.0 | 0.194±0.022 | 0.382±0.034 | 0.283±0.016 | -1.461±0.083 | 0.057±0.001 |
|  | 2-2.5 | 0.185±0.022 | 0.418±0.033 | 0.346±0.014 | -1.464±0.04 | 0.04±0.0 |
|  | 2.5-3.0 | 0.239±0.099 | 0.753±0.105 | 0.391±0.039 | -1.367±0.053 | 0.027±0.0 |
|  | >3 | 0.639±0.065 | 1.082±0.047 | 0.223±0.022 | -1.238±0.041 | 0.008±0.0 |
| PhonBank | 0.5-1 | 0.326±0.065 | 0.391±0.045 | 0.301±0.041 | -1.013±0.087 | 0.035±0.002 |
|  | 1-1.5 | 0.404±0.047 | 0.463±0.021 | 0.446±0.029 | -1.137±0.027 | 0.016±0.0 |
|  | 1.5-2 | 0.891±0.098 | 0.794±0.032 | 0.358±0.042 | -1.234±0.044 | 0.005±0.0 |
|  | 2-2.5 | 1.225±0.136 | 0.877±0.054 | 0.305±0.043 | -1.219±0.046 | 0.002±0.0 |
|  | 2.5-3 | 1.112±0.255 | 0.908±0.1 | 0.38±0.082 | -1.381±0.07 | 0.001±0.0 |
|  | >3 | 1.019±0.371 | 0.857±0.137 | 0.476±0.132 | -1.433±0.087 | 0.001±0.0 |
| *Drosophila* | - | 0.155±0.002 | 0.014±0.0 | 1.1±0.004 | -0.506±0.002 | 0.04±0.001 |
| Zebrafish | - | 0.943±0.054 | 1.33±0.051 | 0.06±0.005 | -0.661±0.052 | 0.0±0.001 |
| Cooking | - | - | - | 0.227±0.029 | -1.133±0.18 | 0.023±0.003 |

### 5.3 Shuffling controls

The speech datasets are organized hierarchically into transcripts, utterances, words, and phonemes allowing us to shuffle the dataset at multiple levels of organization. In the Epic Kitchens, *Drosophila*, and zebrafish datasets no levels of organization were available beyond individual transcripts. To ensure that our MI decay results are a direct result of the sequential organization of each dataset, we performed a control in each dataset in which we shuffled behavioral elements within each individual transcript. In each case, the MI decay is flat confirming that the observed MI decay is a result of sequential organization (Figs S2F, S2E, S10). To ensure that long-range relationships were not due to trivial repetitions of behaviors, we looked in each dataset at MI decay over sequences in which repeated elements were removed. Removing repeats does not qualitatively alter the pattern of long-range relationships between elements (Fig. S4).

### 5.4 Data Availability

The five datasets can be acquired from the TalkBank repository [86], PhonBank repository [85], Berman et al. [56], Damen et al., [99], and Marques et al., [100]. We performed analyses over these transcripts without any modification. Example transcripts for each dataset are displayed in the Supplementary Information. The distribution of sequence lengths of each dataset is shown in Fig. S1. The code necessary for reproducing our results is available on GitHub [117].

## 5.5 Acknowledgements

Work supported by NSF GRF 2017216247 and an Annette Merle-Smith Fellowship to T.S., NIMH training fellowship T32MH020002 and William Orr Dingwall Dissertation Fellowship to A.M., and NIH DC0164081 and DC018055 to T.Q.G.

## 6 Supplementary Materials

**Figure S1:**
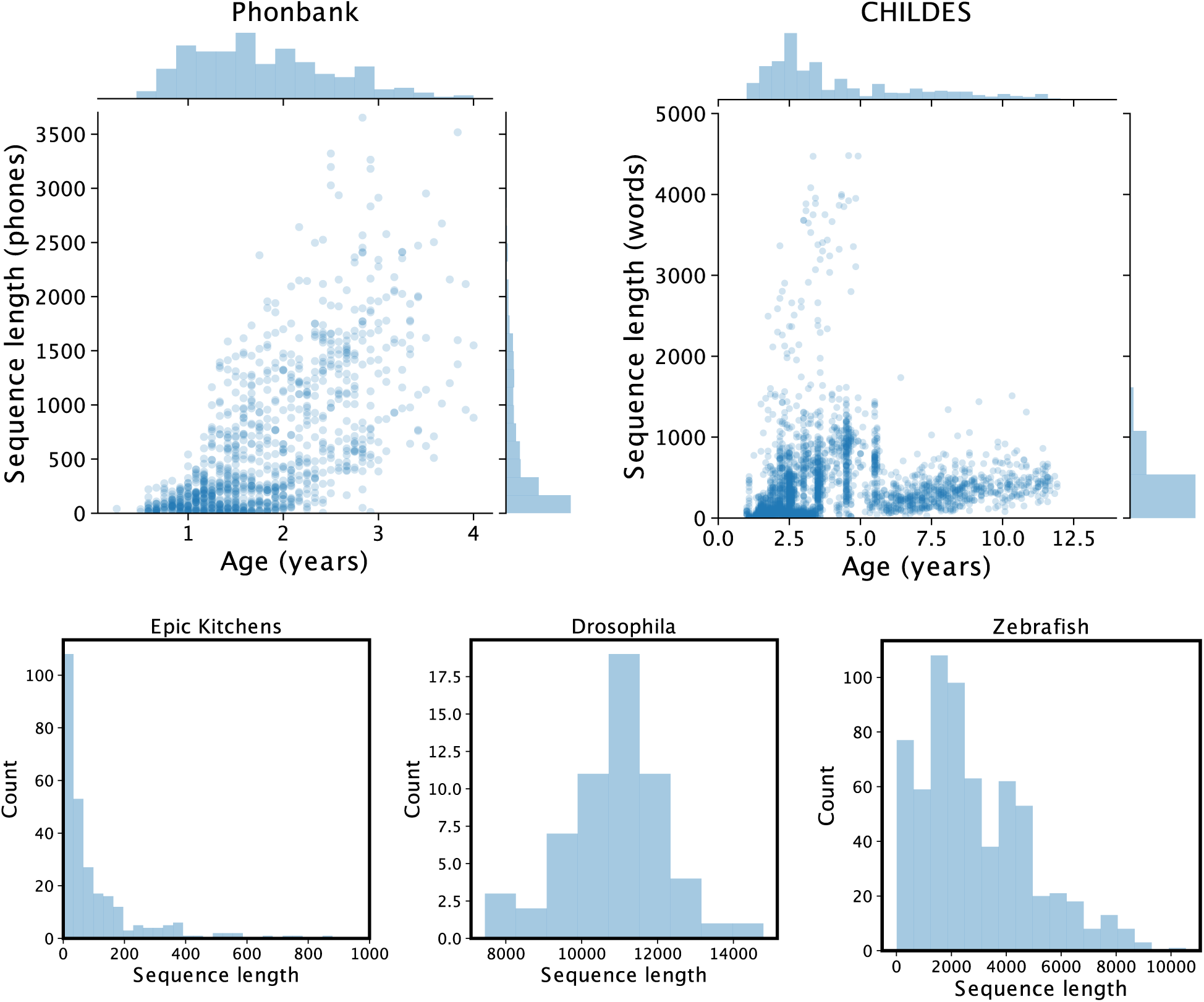
Distribution of sequence lengths for each dataset.

**Figure S2:**
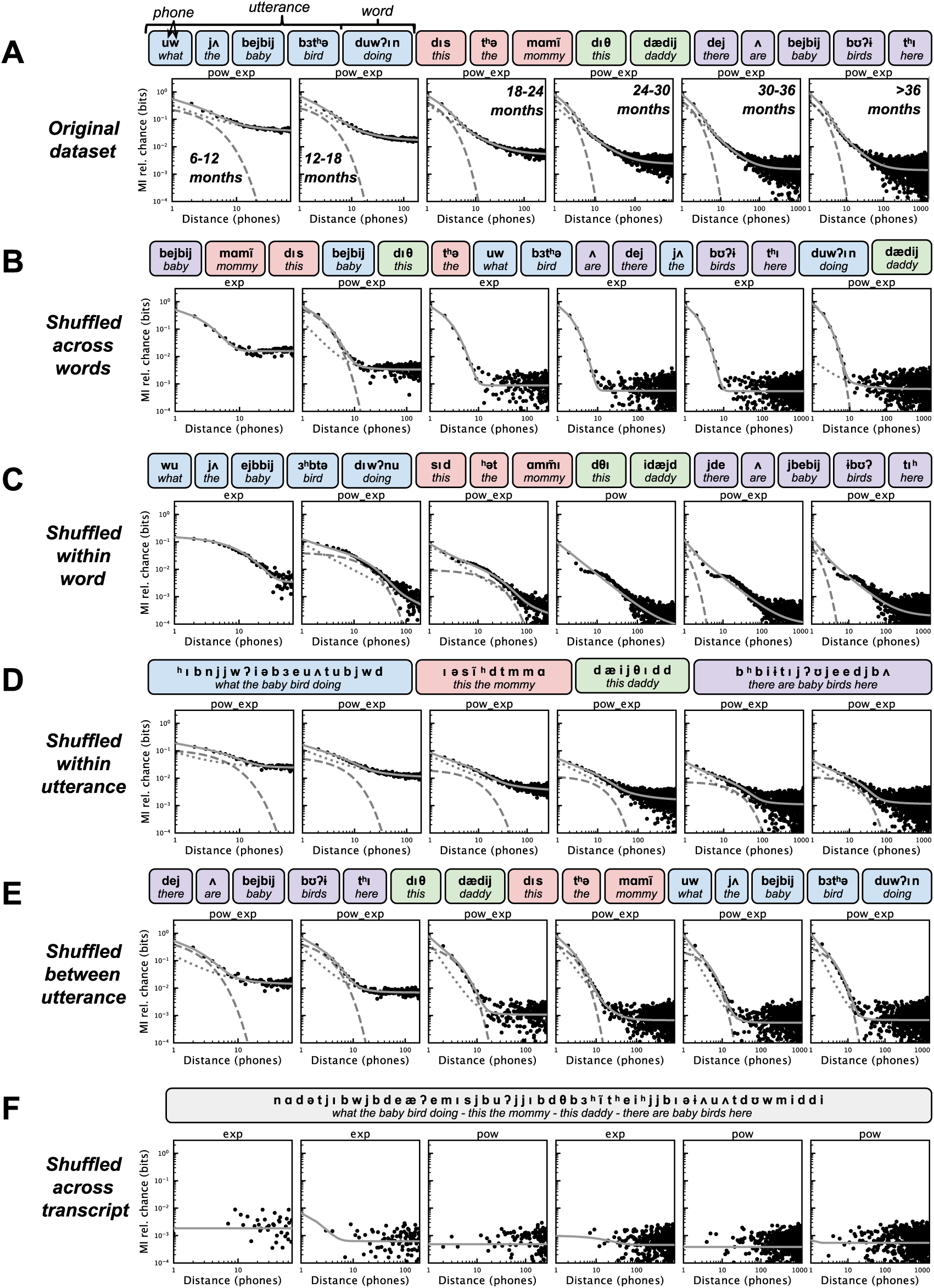
MI decay between phones under different shuffling conditions. (A) MI decay for each age group from the entire dataset, as in Fig. 2A. The sequence above the MI decay shows an example set of utterances of the corpus to illustrate the shuffling conditions. Utterances are grouped by color, words are grouped by rounded rectangles, and phones are displayed in bold above orthographic transcriptions. (B) Words are shuffled within each transcript. (C) Phones are shuffled within words. (D) Phones are shuffled within utterances. (E) Utterances are shuffled within each transcript. (F) Phones are shuffled within each transcript. The best fit model is printed above each plot, and is plotted as grey lies alongside the data and in Fig. 1.

**Figure S3:**
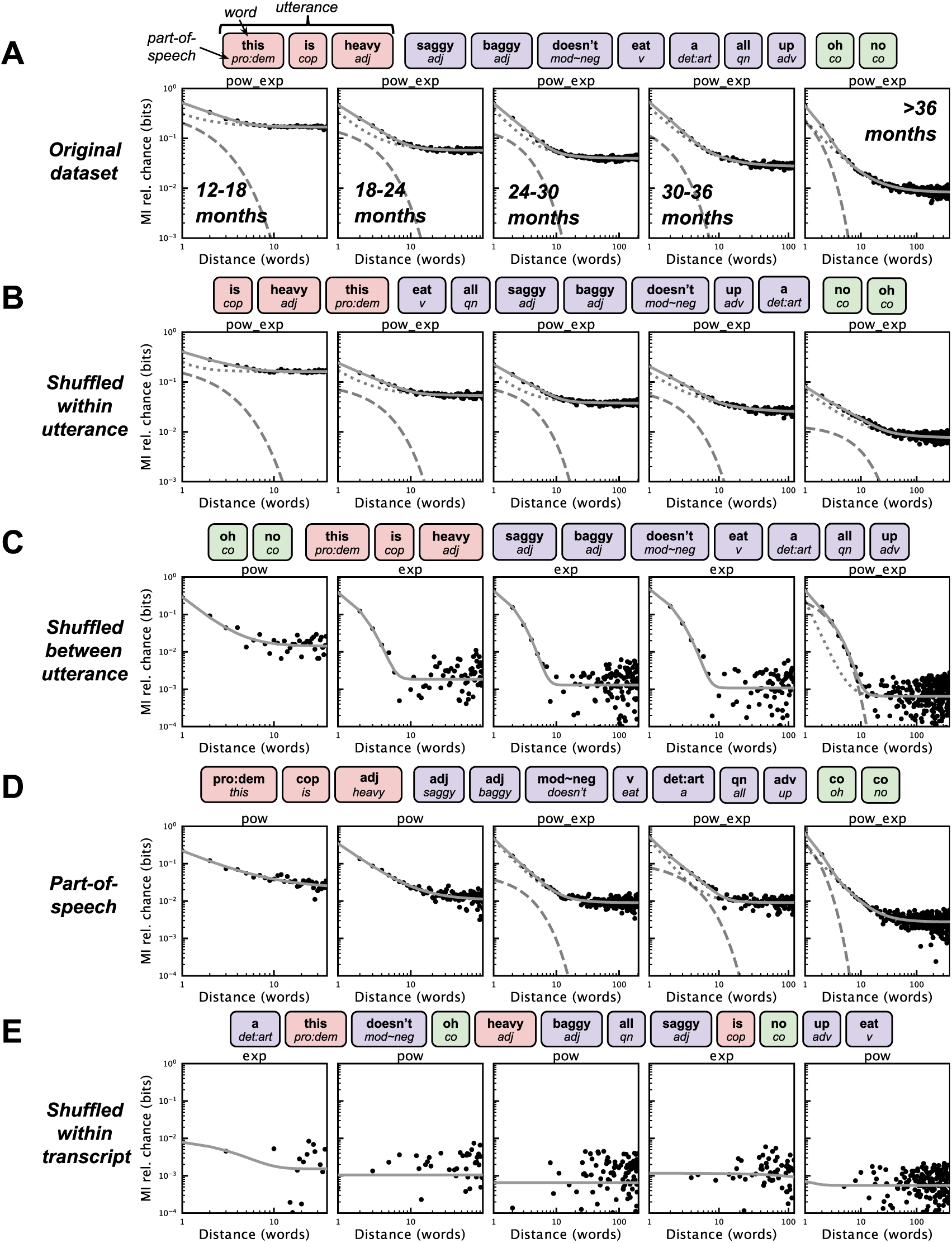
MI decay between words under different shuffling conditions. (A) MI decay for each age group from the entire dataset, as in Fig. 2D. (B) Words are shuffled within each utterance. (C) Utterances are shuffled within each transcript. (D) MI is calculated over part-of-speech transcriptions of words. (E) Words are shuffled within each transcript. (F) Words are shuffled within each transcript. The best fit model is printed above each plot, and is plotted as grey lies alongside the data and in Fig. 1.

**Figure S4:**
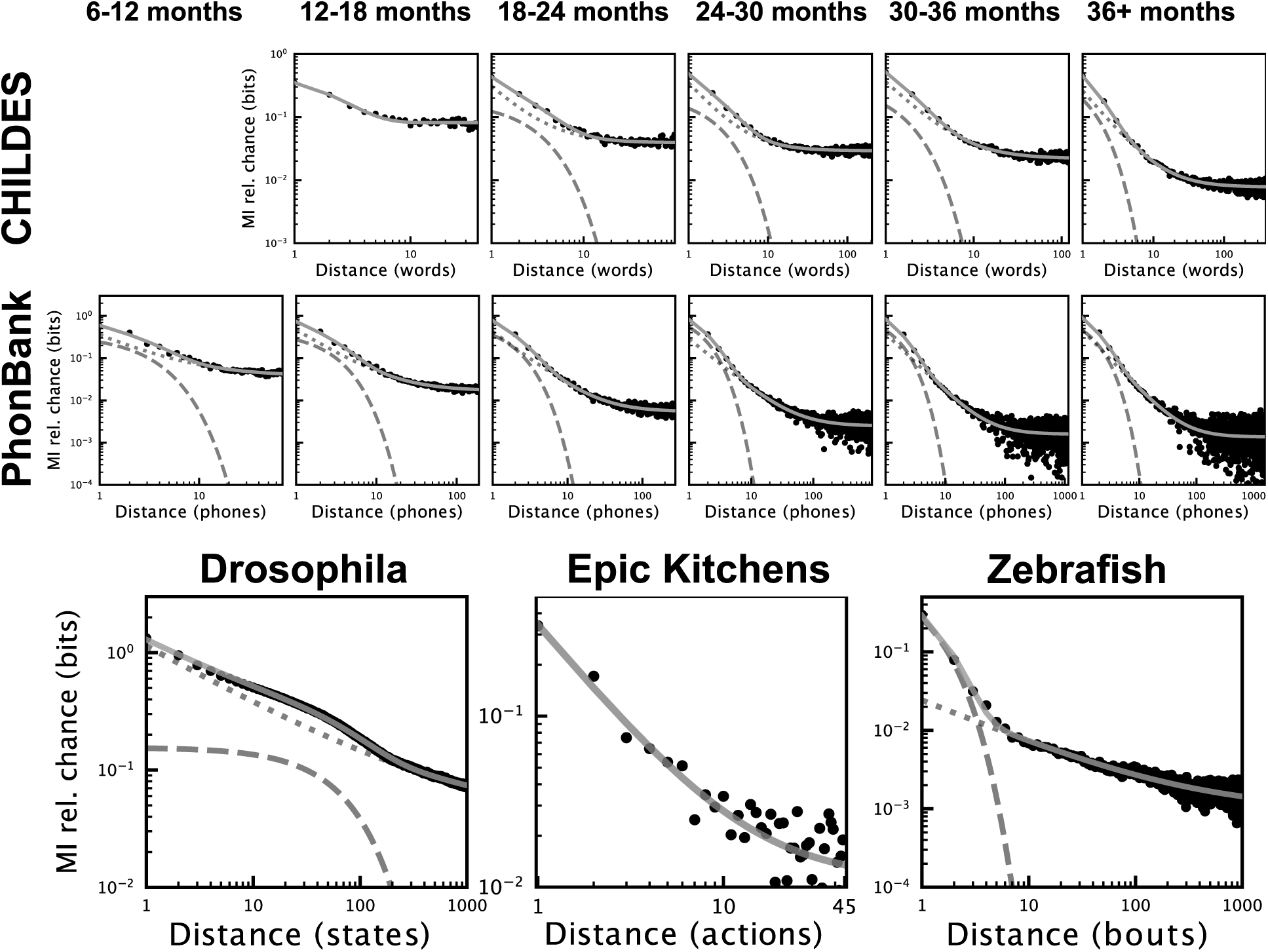
MI decay with repeated elements removed across each dataset.

**Figure S5:**
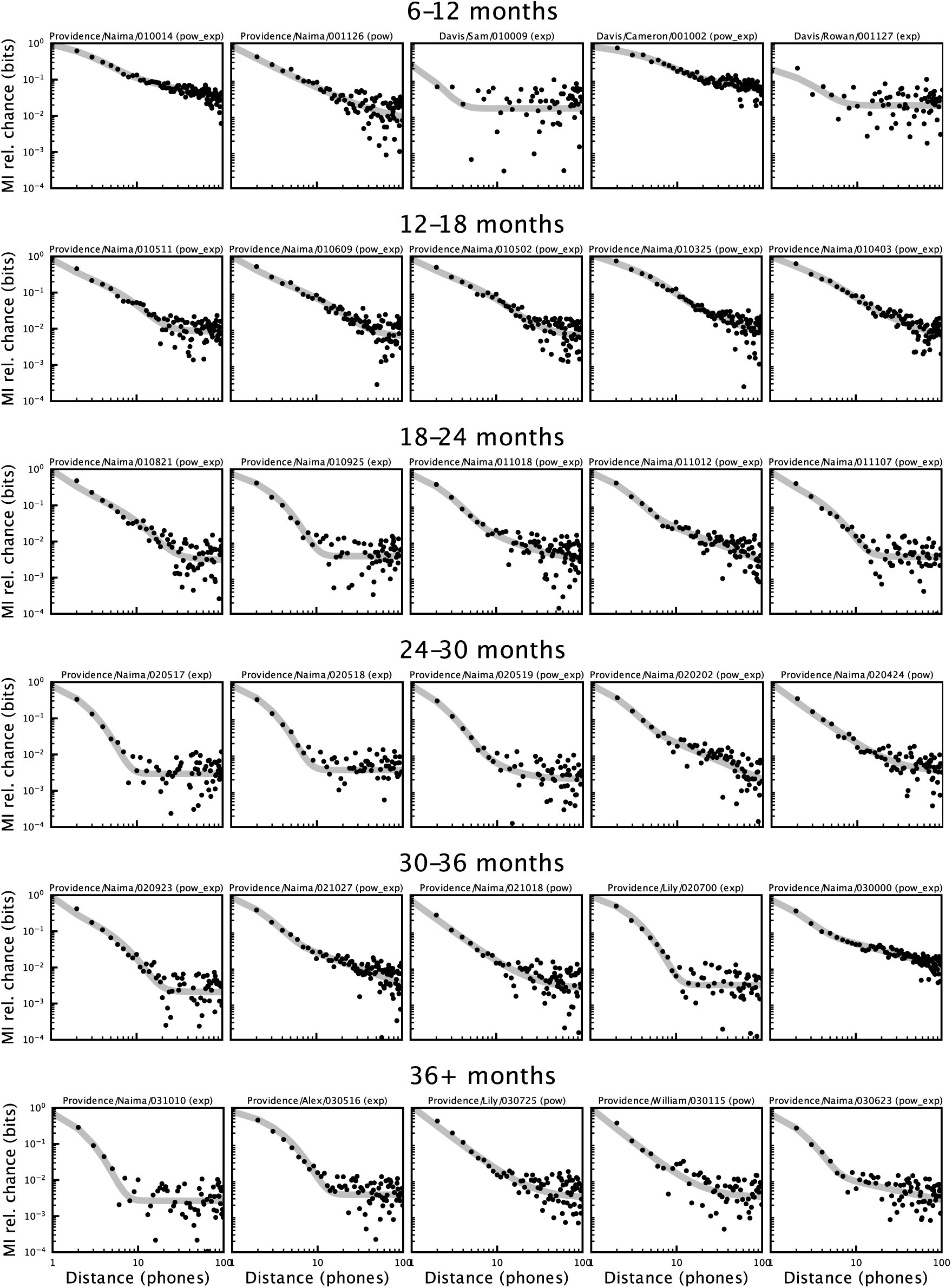
MI decay and best fit model of five largest transcripts for each age group across PhonBank. Transcript identity and best fit model are displayed above each plot.

**Figure S6:**
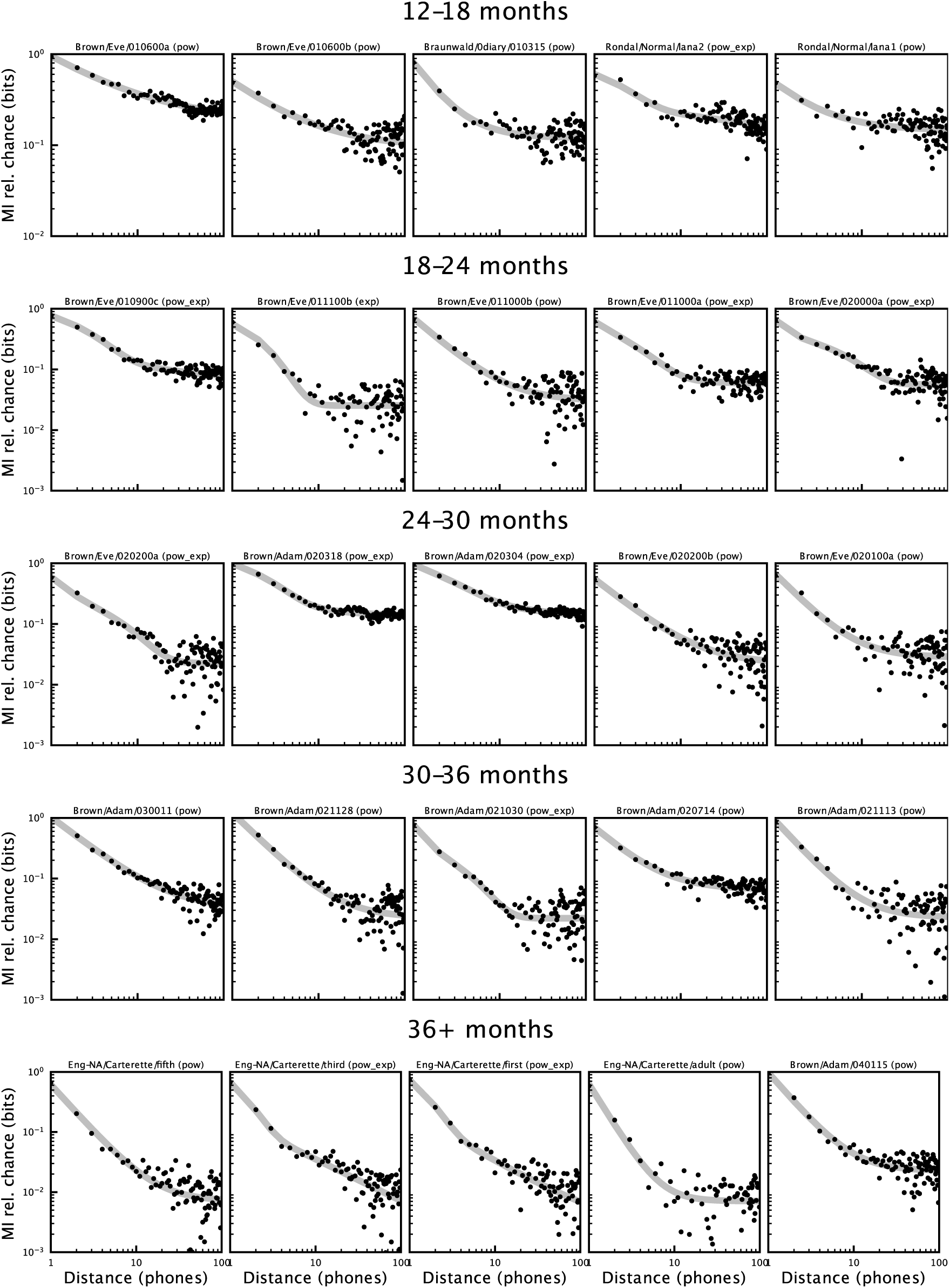
MI decay and best fit model of five largest transcripts for each age group across CHILDES. Transcript identity and best fit model are displayed above each plot.

**Figure S7:**
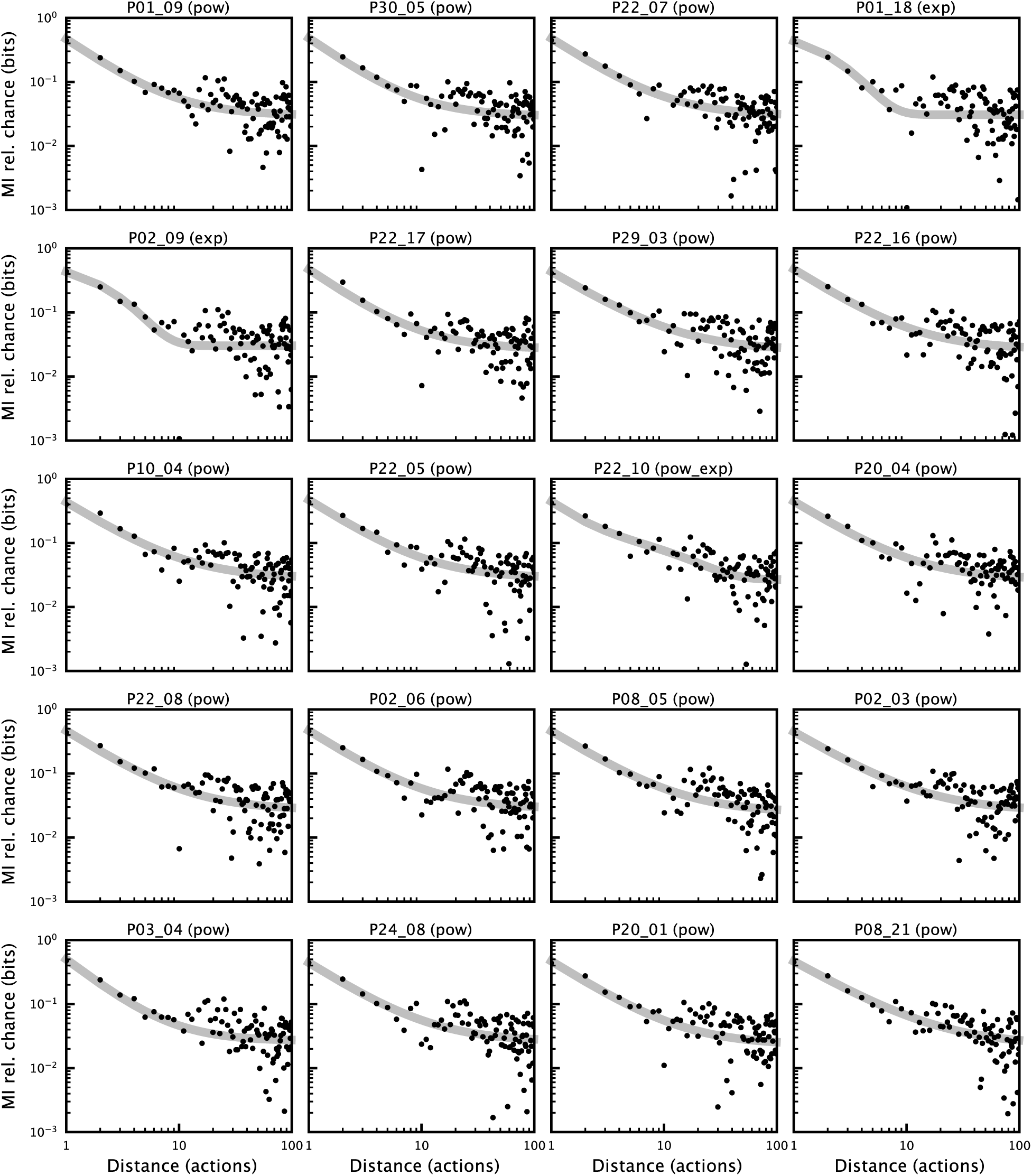
MI decay over the 20 longest Epic kitchens cooking sequences. Transcript identity and best fit model are displayed above each plot.

**Figure S8:**
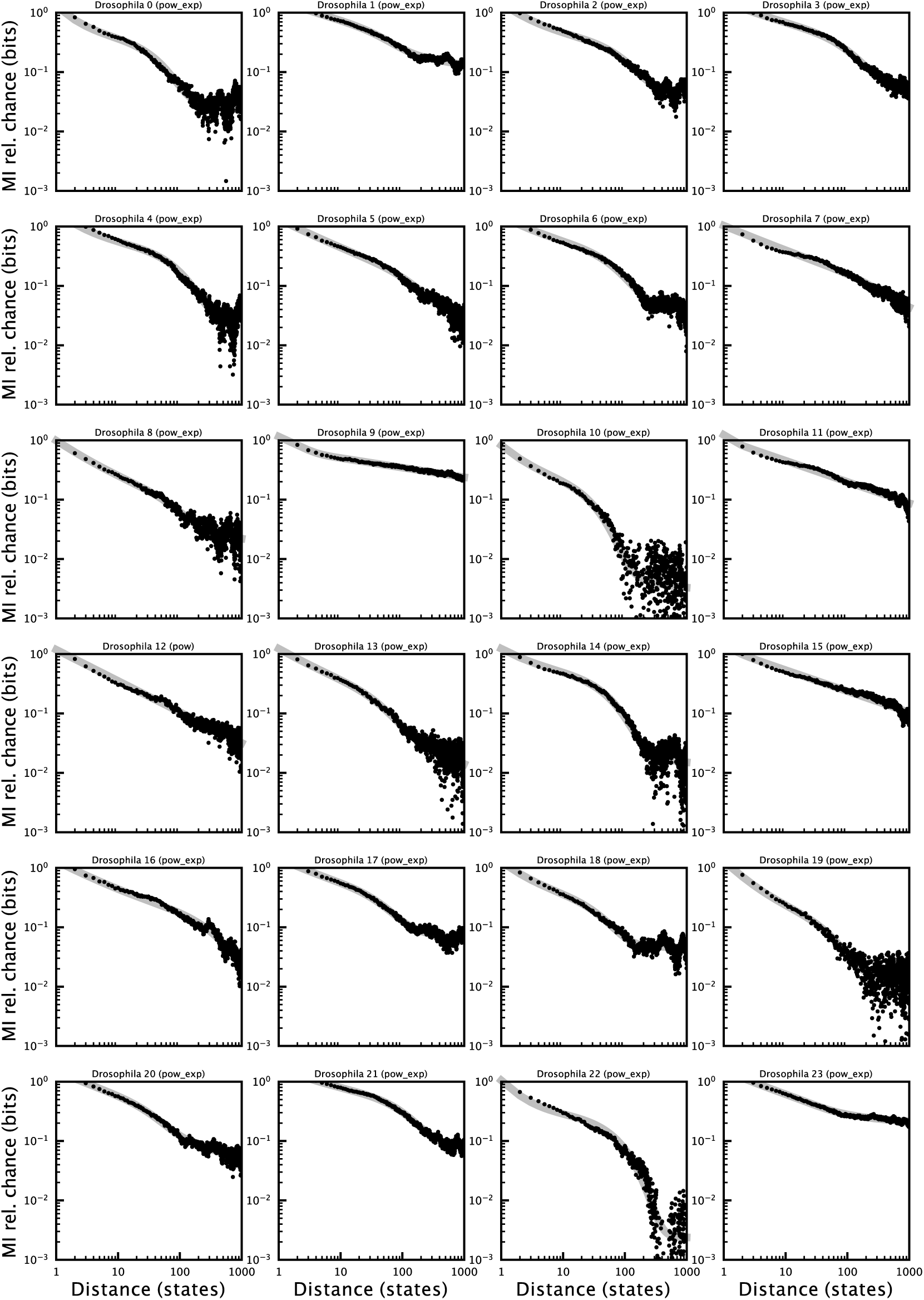
MI decay of example individual Drosophila behavioral sequences over one hour. Transcript identity and best fit model are displayed above each plot.

**Figure S9:**
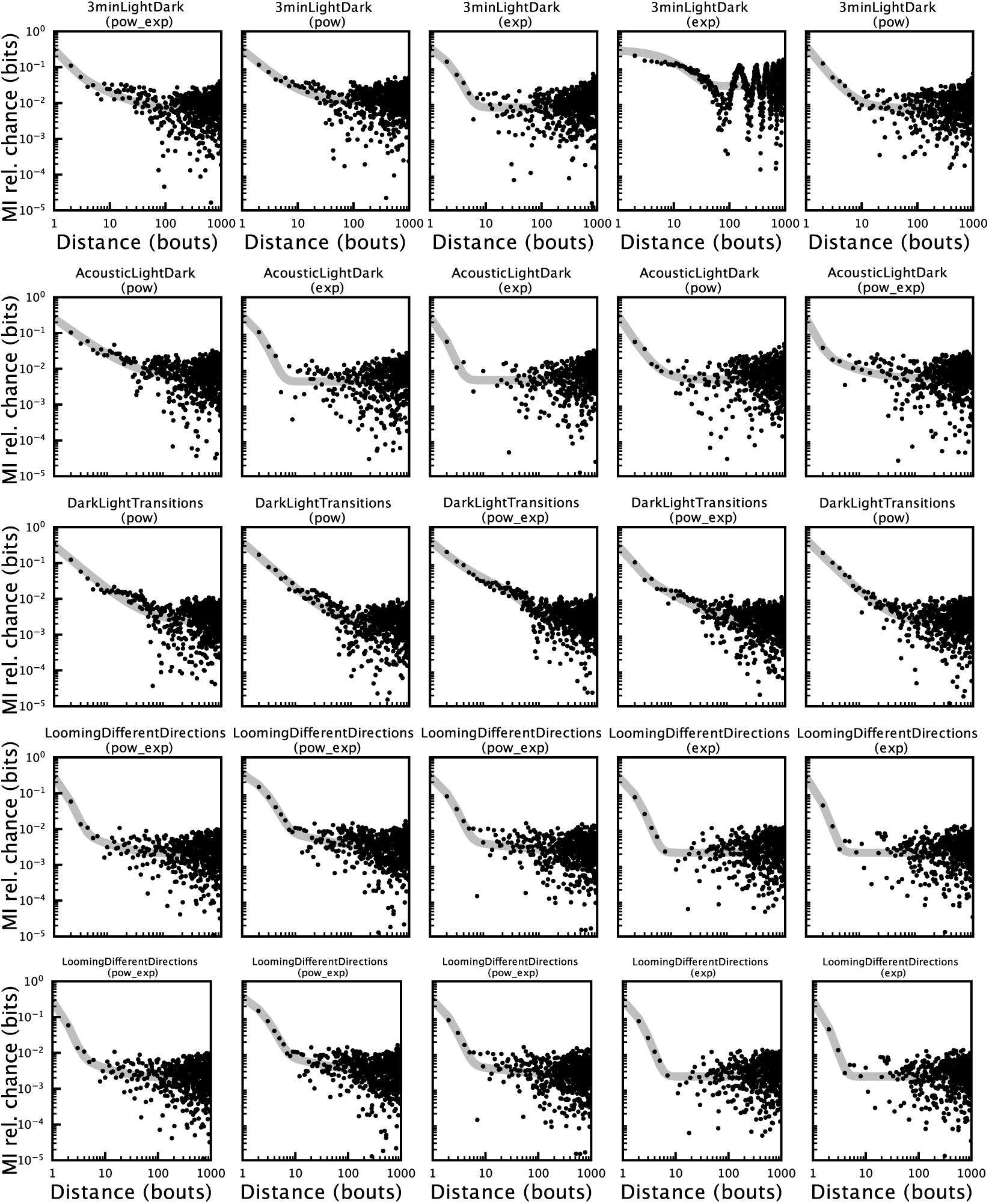
MI decay of several individual Zebrafish behavioral sequences. Each plot corresponds to the continuous behavior of a single Zebrafish. Each row corresponds to a different behavioral setting. The behavioral setting is written above the plot alongside the best fit model.

**Figure S10:**
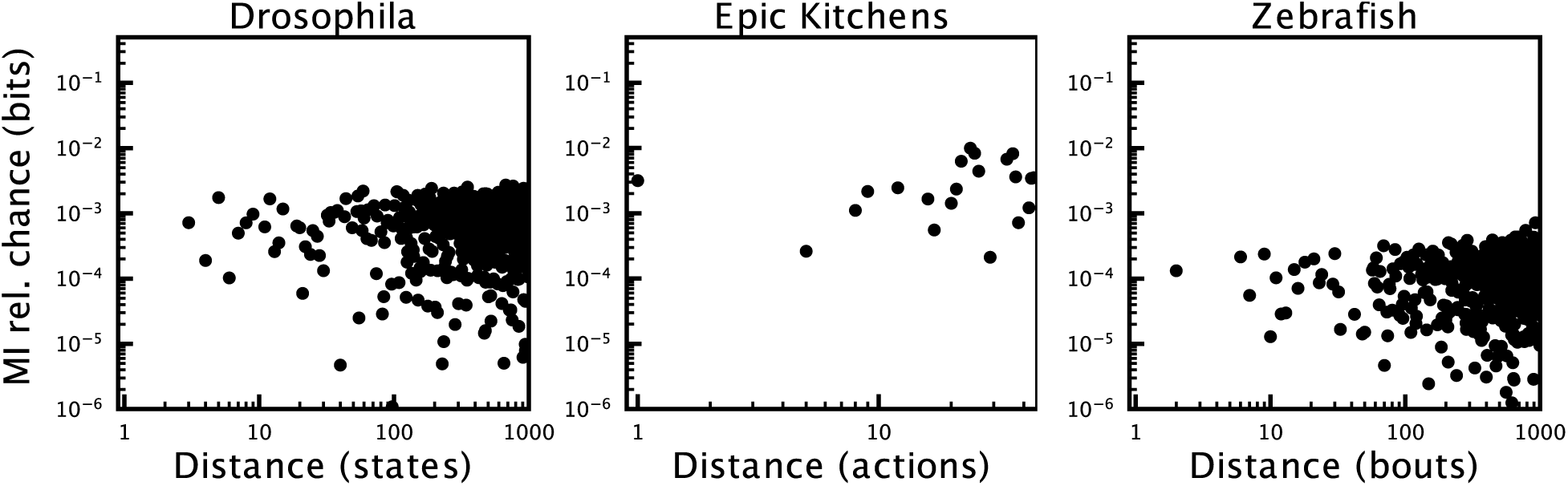
MI decay of shuffled sequences for Drosophila, Zebrafish, and Epic Kitchens datasets. No information decay is seen between elements of any sequence.

## 7 Example sequences from datasets

### 7.1 PhonBank

A random sample of the transcripts used in this manuscript at different ages. Each line corresponds to an utterance and each utterance is followed by an orthographic representation in parentheses. ‘xxx’ in orthographic transcription refers to unintelligible speech and ‘yyy’ refers to phonological coding. The meanings of other coding symbols such as ‘@’ and ‘&’ used in orthographic representations can be found in the TalkBank manuals for PhonBank and CHILDES.

#### 7.1.1 Davis/Nate/001105.xml 11 months

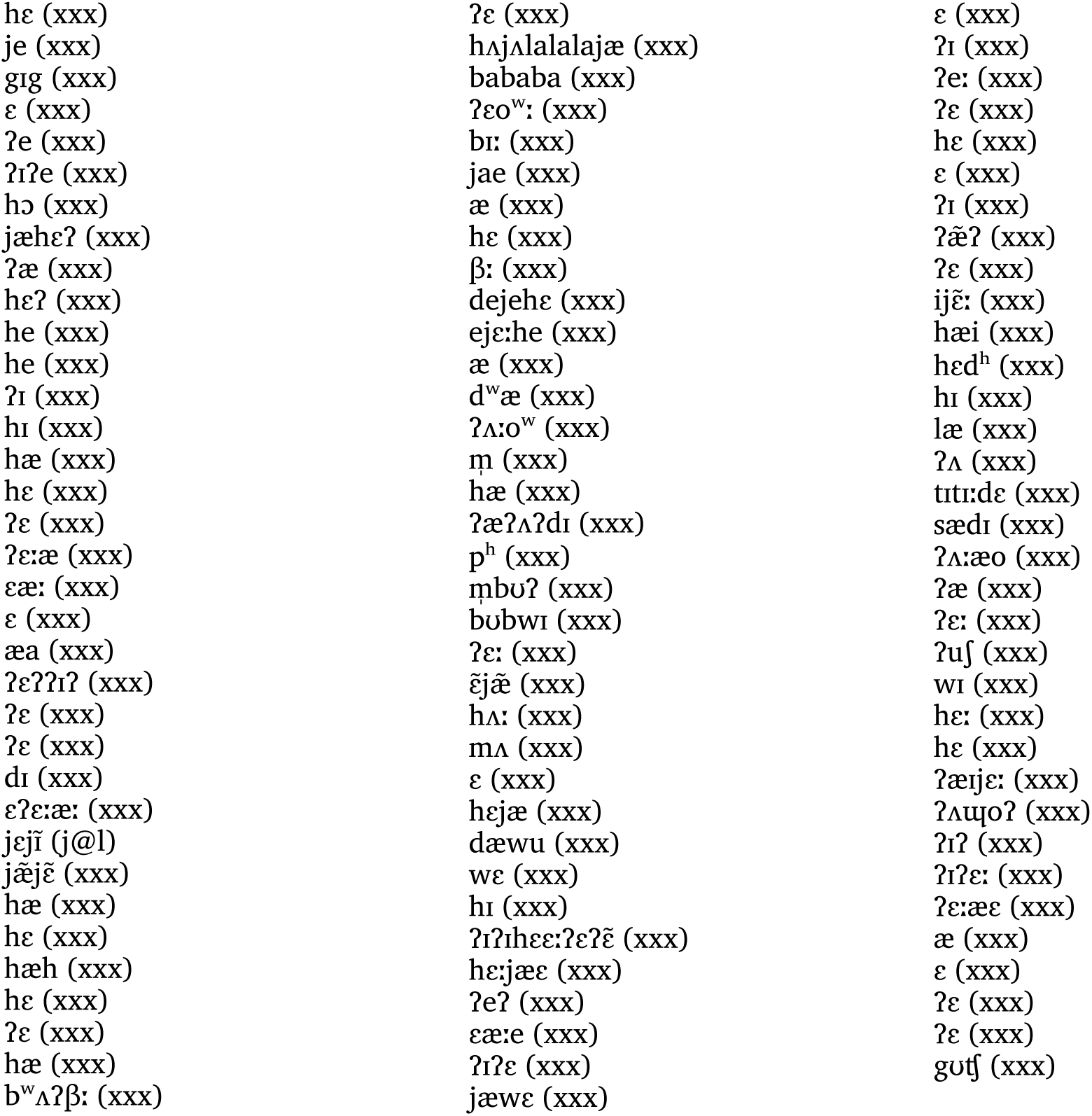

#### 7.1.2 Providence/William/011115.xml 23 months

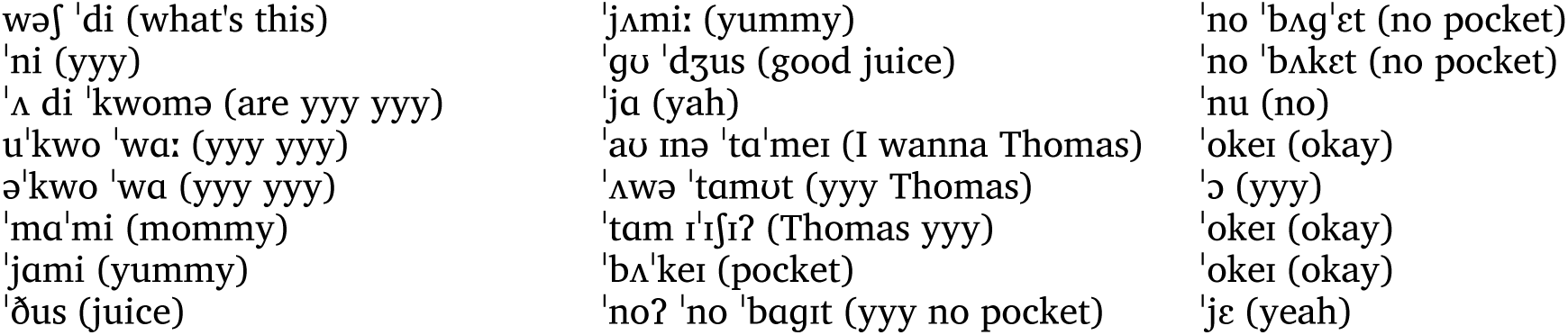
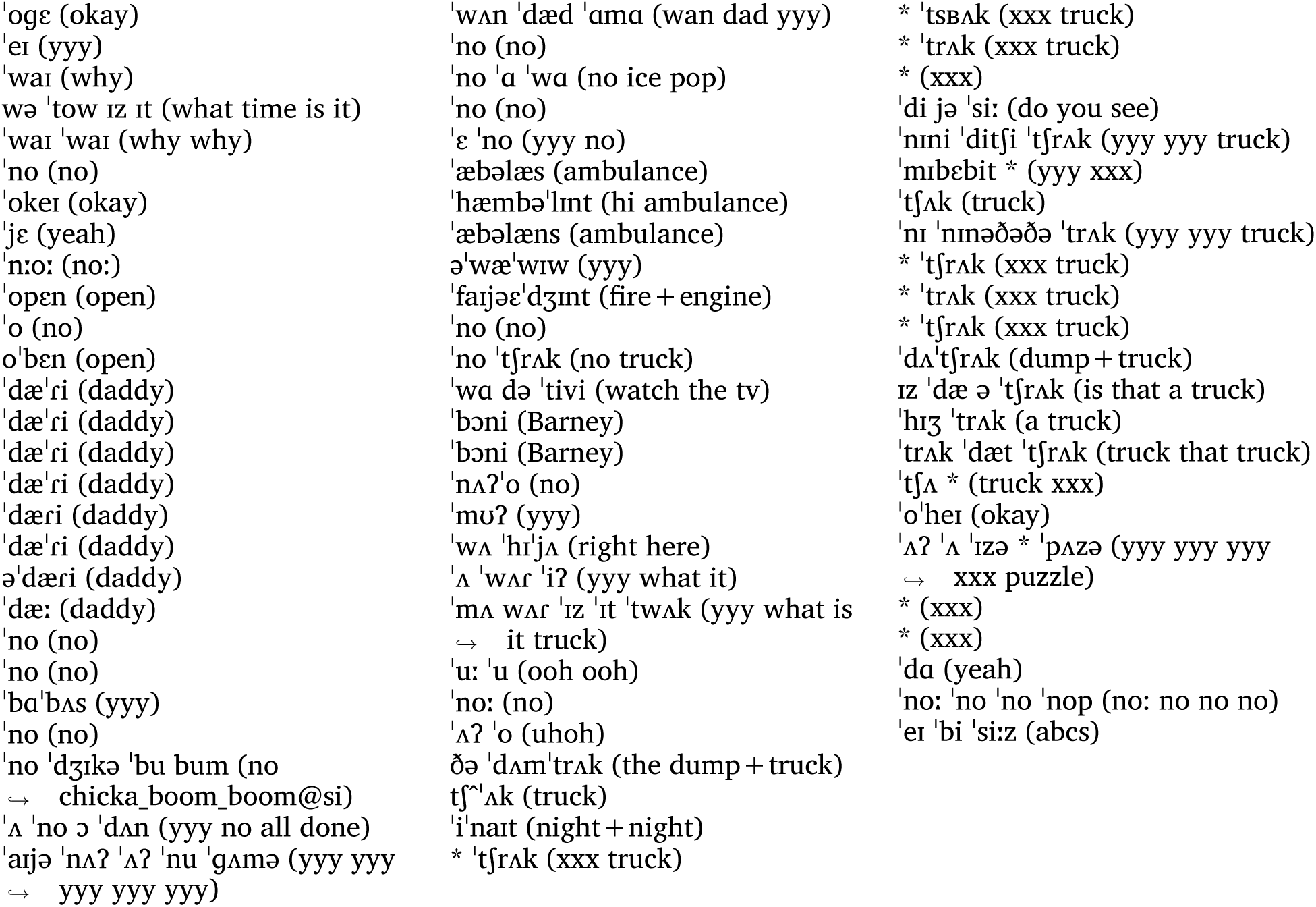

#### 7.1.3 Goad/Julia/20510.xml 29 months

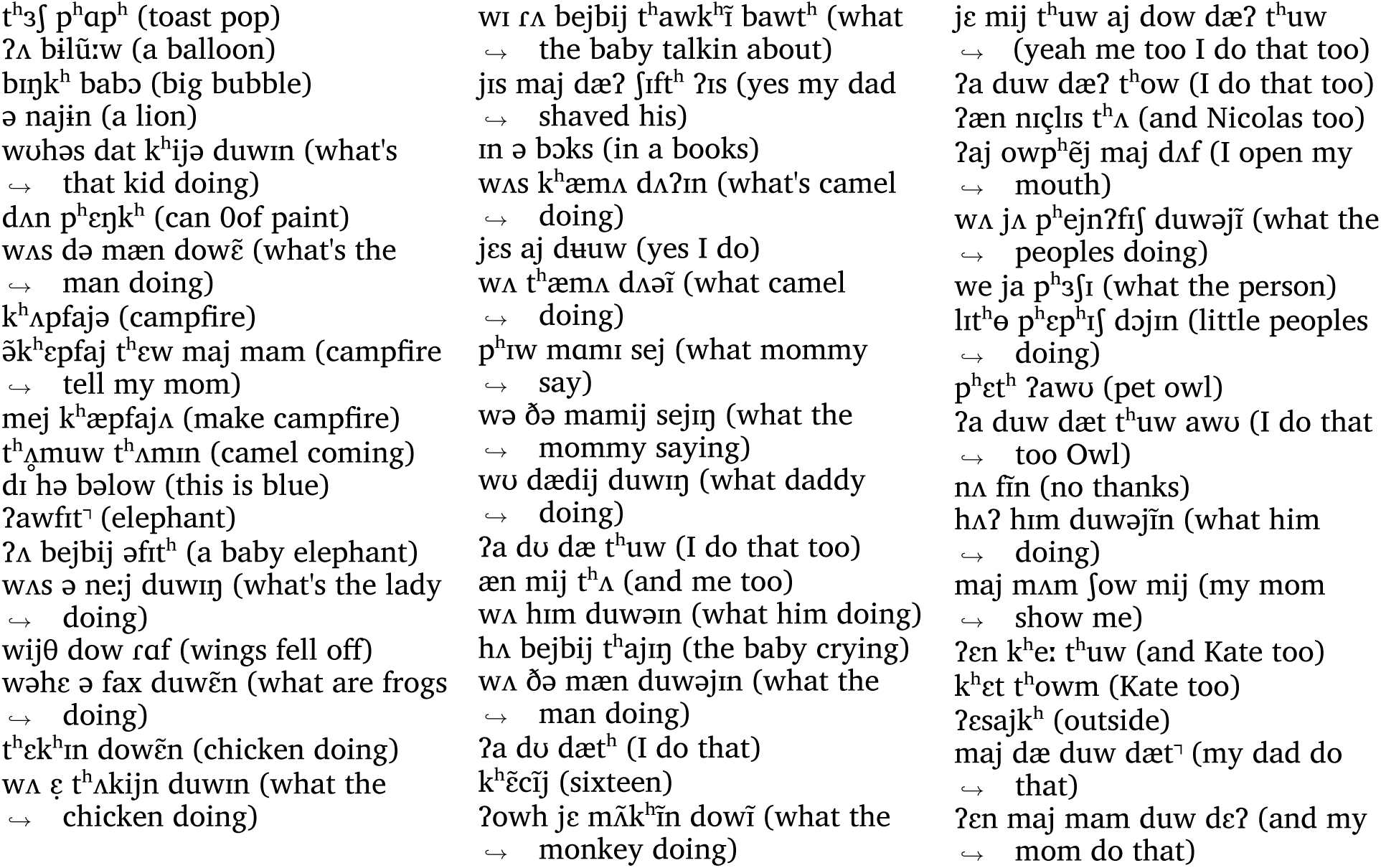
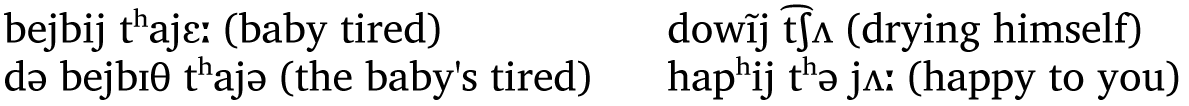

#### 7.1.4 Providence/Alex/021122.xml 36 months

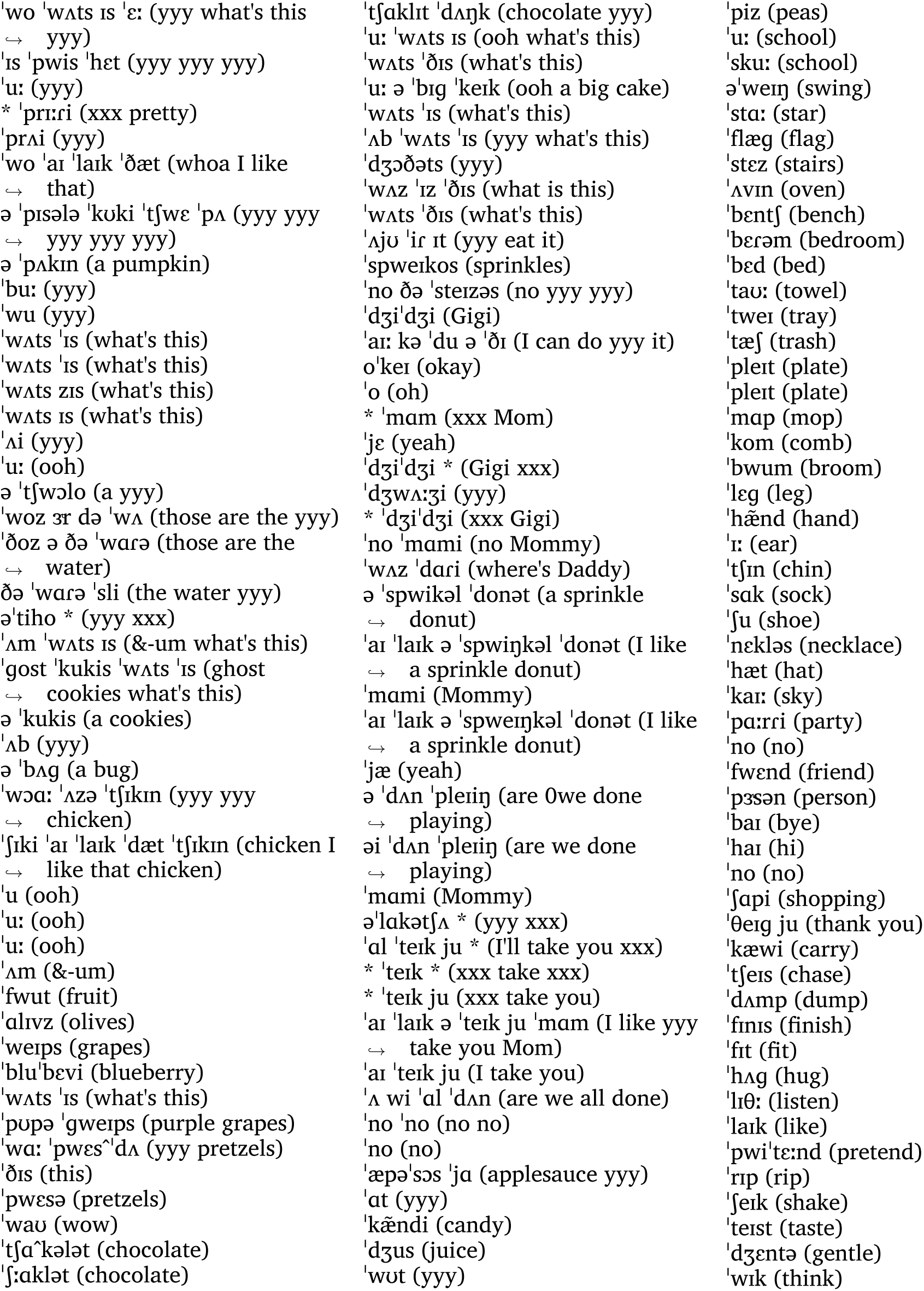
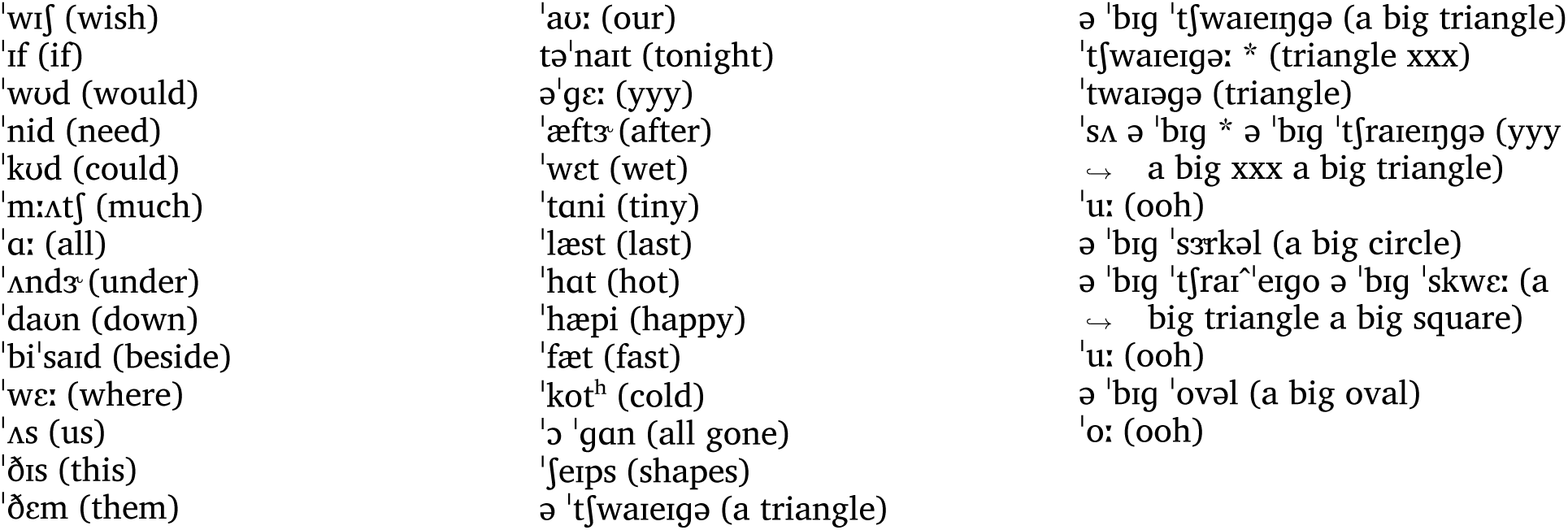

### 7.2 CHILDES

A random sample of the transcripts used in this manuscript at different ages. Each line corresponds to an utterance and each utterance is followed by transcribed part-of-speech tags.

#### 7.2.1 Eng-NA/Braunwald/010511.xml 17 months

|  |  |  |
| --- | --- | --- |
| night_night (co) | yeah (co) | she lives next door to us (pro:sub |
| night_night (co) | on (adv) | ↪ v adj n prep pro:obj) |
| here (adv) | Cee (n:prop) | bow (on) |
| it is night_night (pro:per 0cop n) | spider (n) | recorder (n) |
| Daddy (n:prop) | Cee (n:prop) | cookie (n) |
| spiders (n) | down (adv) | no (co) |
| oh (co) | byebye (co) | Deedee (n:prop) |
| me Dwww (pro:obj n:prop) | car (n) | here (adv) |
| on (adv) | car (n) | cookie (n) |
| on (unk) | there (adv) | that that door (det:dem n) |
| no (co) | byebye (co) | that tata (comp chi) |
| buttons (unk) | car (n) | nose (n) |
| uh () | car (n) | eye (n) |
| down (adv) | baby (n) | ear (n) |
| water (n) | night_night (co) | Laura (n:prop) |
| water (n) | night_night (co) | toe (n) |
| there (adv) | Cee (n:prop) | tickle (n) |
| dance there (unk adv) | cookie (unk) | toe (n) |
| ahhah (co) | spoon (n) | ah (co) |
| on (adv) | oh (co) | uh () |
| don't (mod~neg) | down (unk) | toe (n) |
| give (v) | down (unk) | recorder (unk) |
| I want (pro:sub v) | there (adv) | toe (n) |
| Daddy (n:prop) | recorder (n) | ah (co) |
| dance (n) | aya (bab) | toe (n) |
| on (adv) | door (n) | my toe (det:poss n) |
| I want that that that that | key (n) | toe (n) |
| ↪ (0pro:sub v pro:dem) | byebye (co) | where (pro:rel) |
| eh () | car (n) | here (adv) |
| go in there (v prep n) | kitty (n) | no (co) |
| uhoh (co) | outside (adv) | there (adv) |
| uhoh (co) | bow (on) |  |
| uhoh (co) | bye (co) |  |
| yeah (co) | byebye (co) |  |
| thank you (v pro:per) | bow (on) |  |
| thank you (v pro:per) | bow (on) |  |

#### 7.2.2 Brown/Adam/020801.xml 32 months

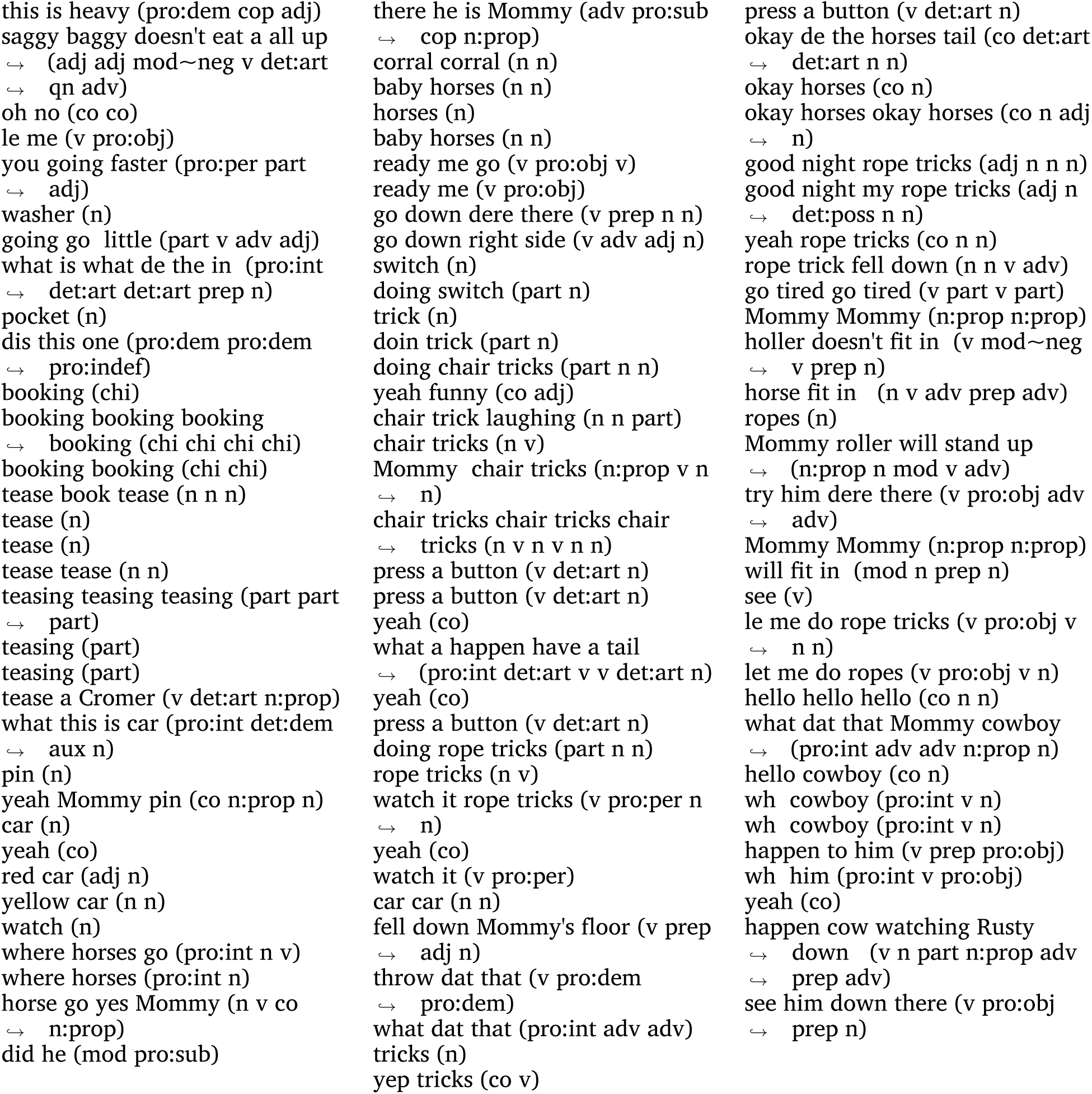

#### 7.2.3 Eng-NA/Carterette/first.xml 72 months

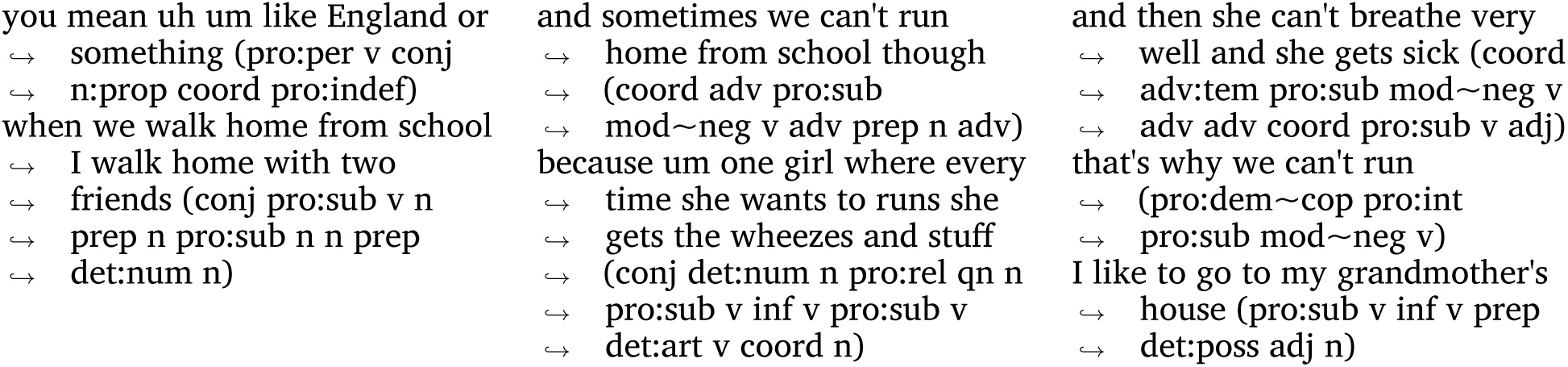
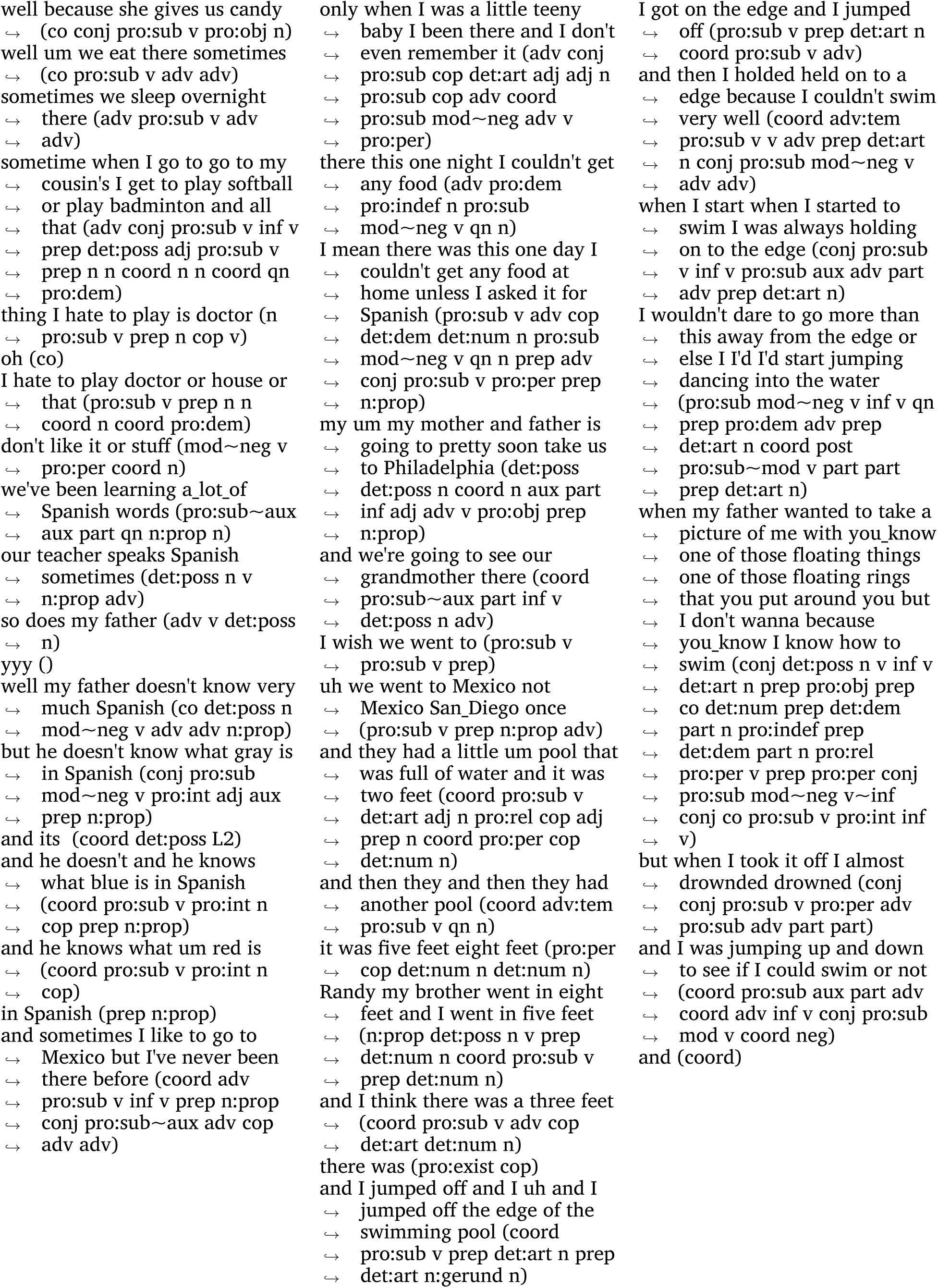
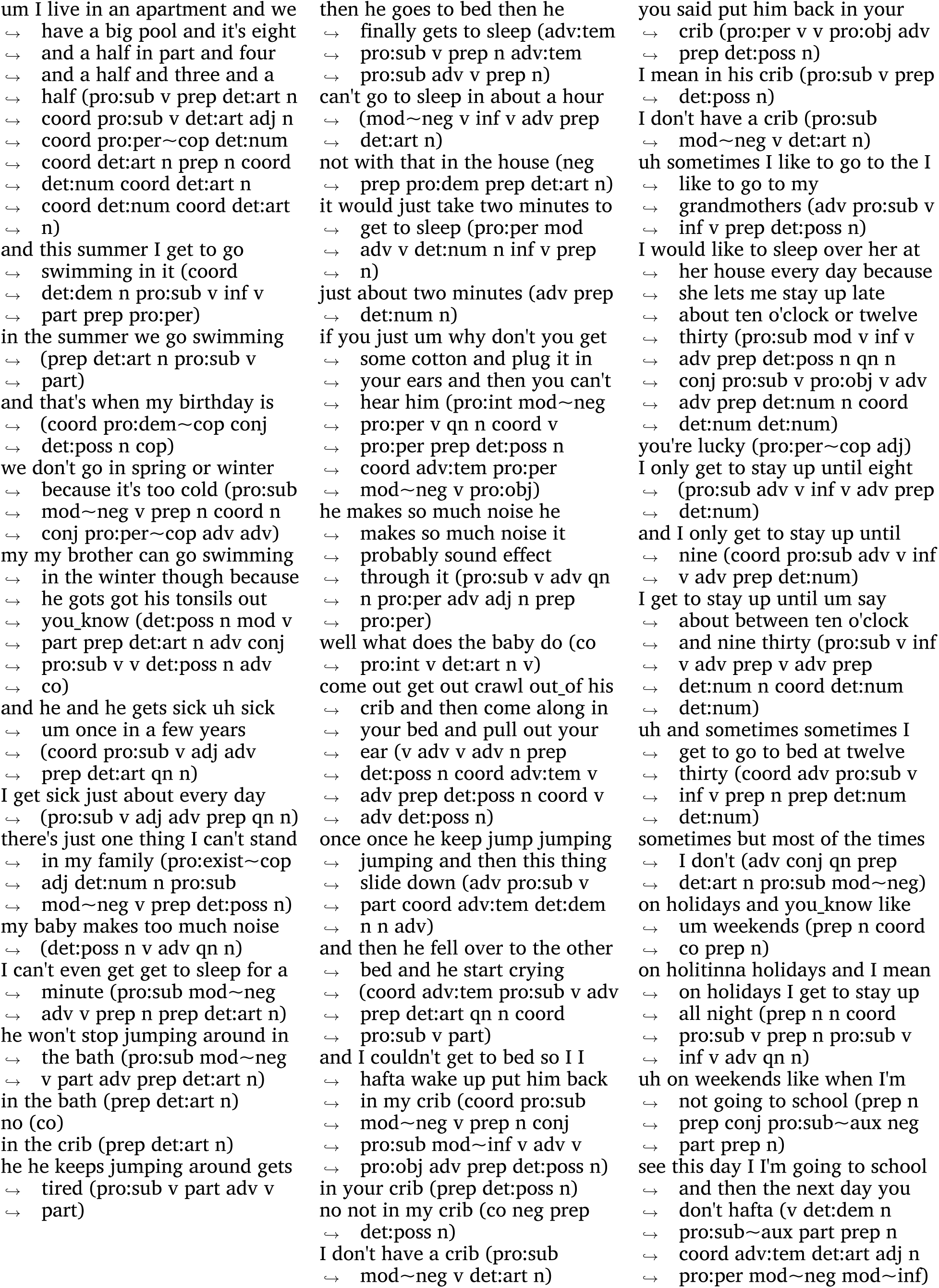
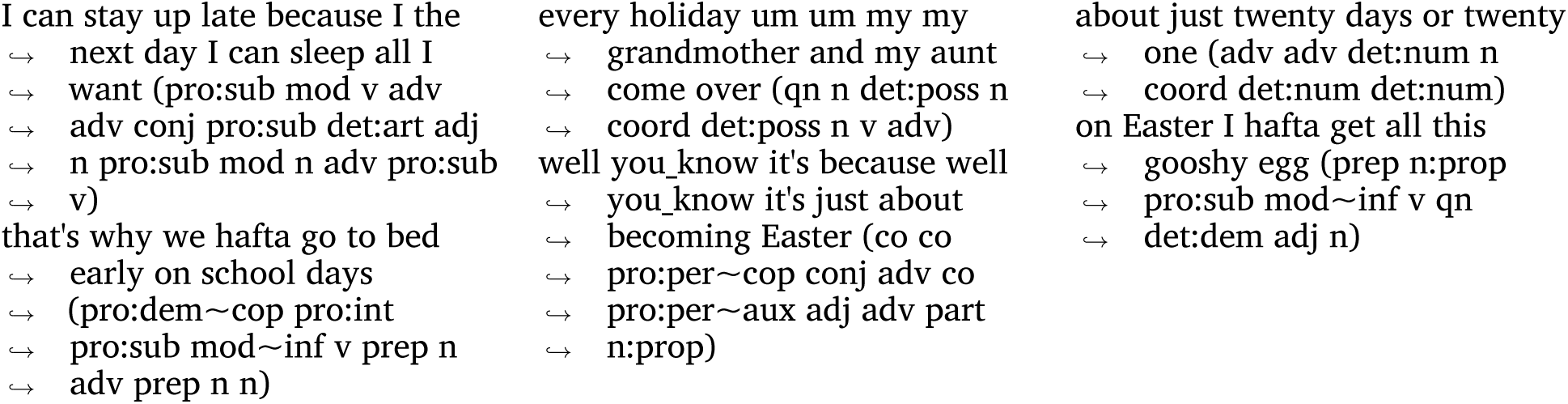

### 7.3 Drosophila

One hour of behavioral state transitions from a single example *Drosophila*. There are 117 unique behavior states. Behavioral states do not have names but belong to broad categories (Posterior, Side Legs, Anterior, Locomotion, Idle, Slow).

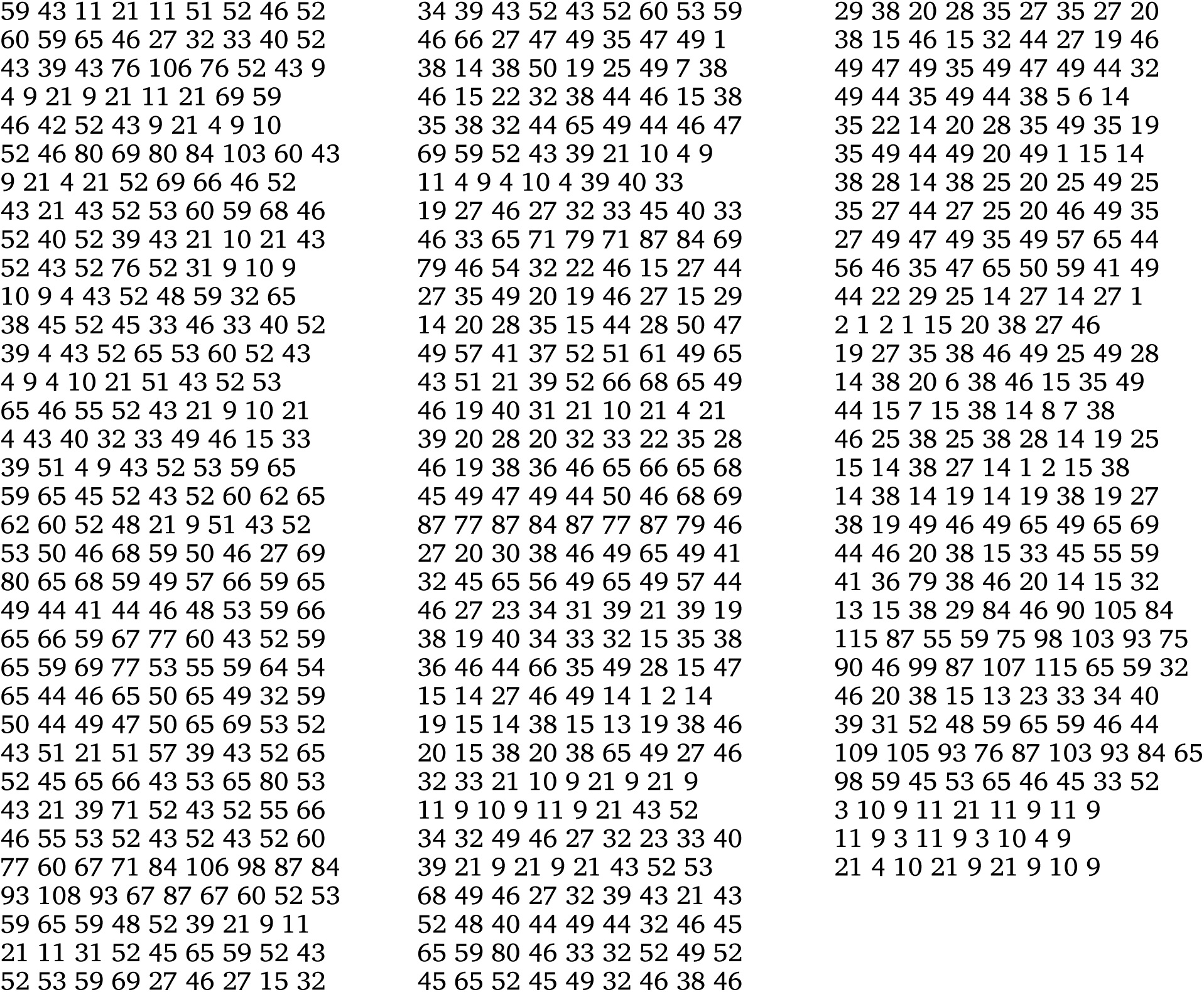

### 7.4 Zebrafish

Behavioral states for zebrafish. Several behavioral contexts are used in this dataset. The example behavioral sequence shown below is acquired during a phototaxis paradigm (SCS: Short Capture Swims; LCS: Long Capture Swims; BS: Burst type forward Swim with high tail-beat frequency; SLC: Fast C-start escape Swims; RT: Routine Turns; LLC: Long Latency C-starts; AS: Approach Swims; SAT: Spot Avoidance Turn; HAT: High Angle Turn).

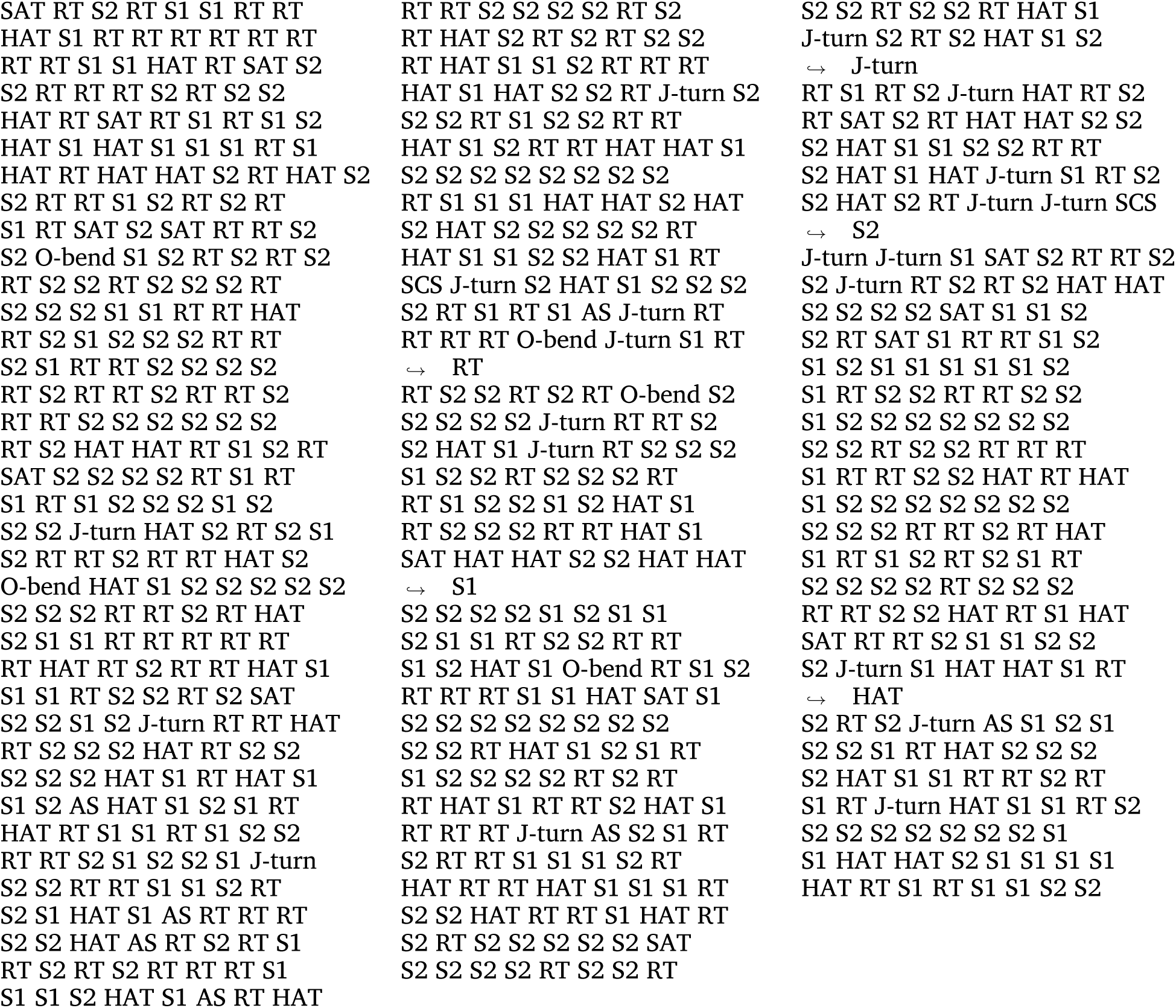

### 7.5 Epic Kitchens

Each transcript in Epic Kitchens contains a sequence of behaviors consisting of an action and object. One example sequence is shown below.

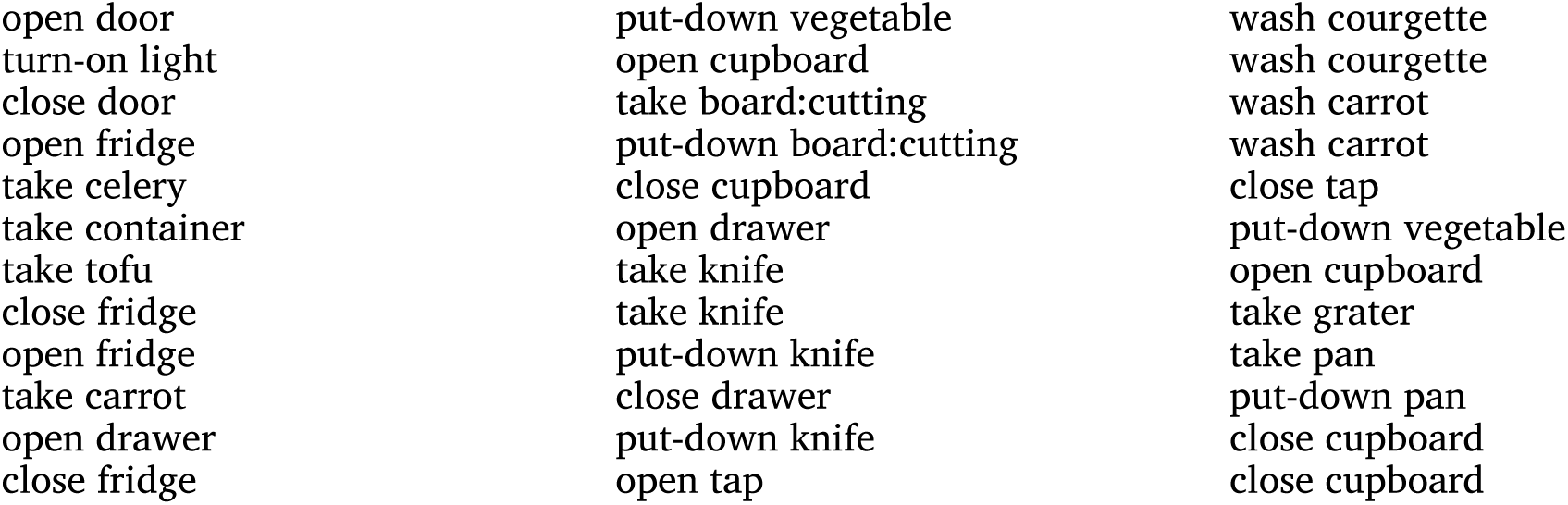
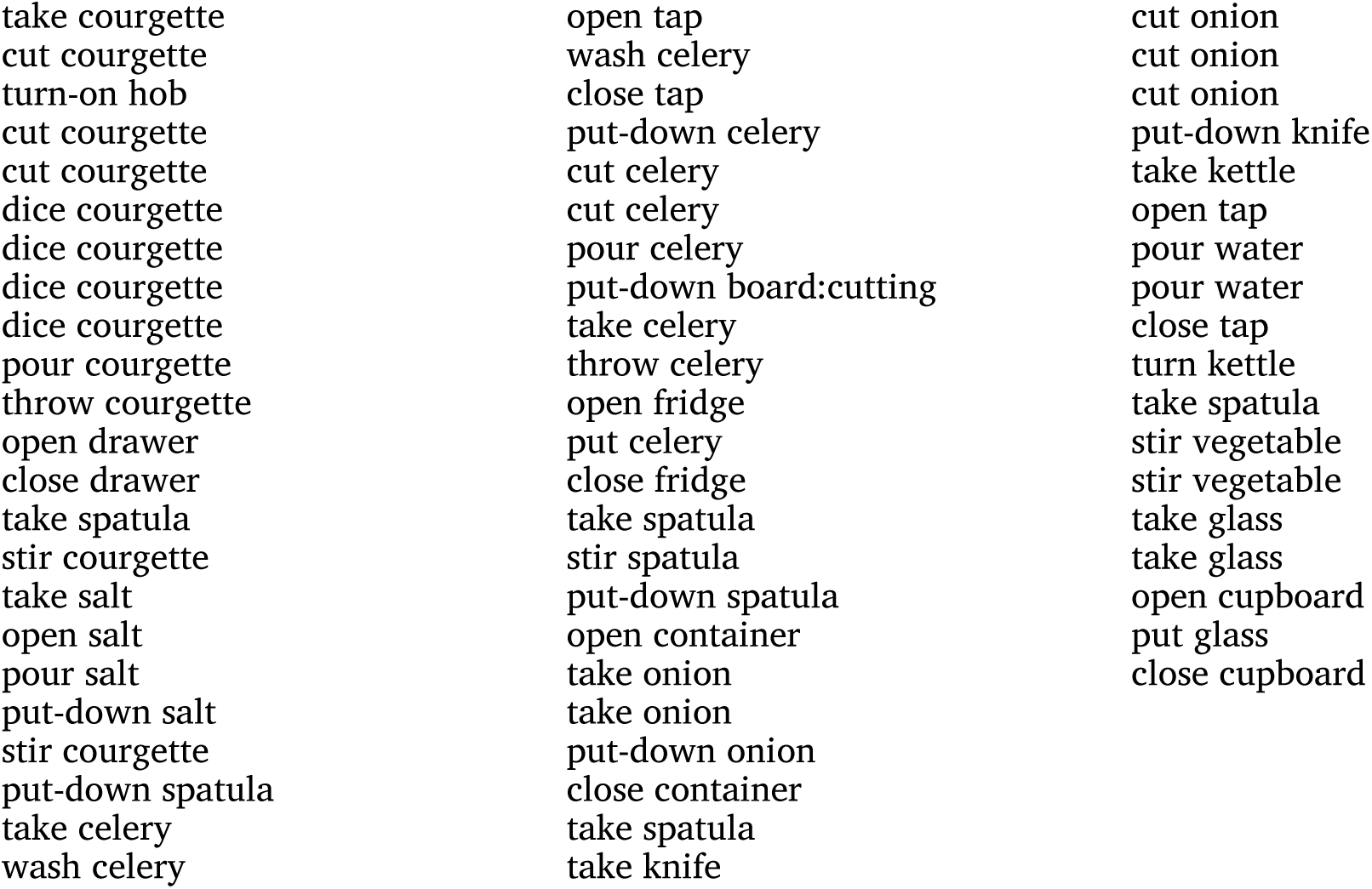

